# *Caenorhabditis elegans* avoids *Todstoff*, a novel nociceptive necrotaxis cue

**DOI:** 10.64898/2026.02.06.704436

**Authors:** Erik Toraason, Rachel Kaletsky, Jonathan Z. Huang, Vanessa Y. Ying, Xuanjia Ye, Edoardo Longarini, Alden Weiner, Waasae Ayyaz, Pablo Ramirez-Garcia, Tom W. Muir, Mohammad R. Seyedsayamdost, Coleen T. Murphy

## Abstract

To survive in hazardous environments, organisms must navigate and avoid myriad threats. Many species across phyla detect warning cues in the remains of dead or injured conspecifics, allowing them to enact defensive strategies and avoid a similar fate. Here, we demonstrate that *Caenorhabditis elegans* senses a novel aversive cue (which we term ‘*Todstoff*’, or ‘death substance’) present in the remains of dead worms that induces living conspecifics to perform negative necrotaxis behaviors. Todstoff is distinct from previously-identified social signals in *C. elegans*, including ascaroside and alarm pheromones, and our biochemical analysis has revealed that Todstoff may be a protein-associated amino acid derivative less than 1.35 kDa in size. We determined that Todstoff is sensed by the ASH polymodal nociceptive neurons, and further, that the necrotaxis signal is transduced via the activity of G-protein coupled receptor (GPCR) signaling, the Gi/o-like protein ODR-3, TRPV channels OSM-9 and OCR-2, and glutamatergic synaptic transmission to downstream AIB interneurons. Taken together, our work illuminates a post-mortem inter-animal chemical signaling pathway that promotes death avoidance, and thus, survival.

## Introduction

Organisms encounter diverse threats in their environment. In addition to intrinsic mechanisms to tolerate exogenous assault, such as stress and immune pathways, many species have evolved the capacity to recognize and avoid deceased conspecifics^1–8^. The logic to this strategy is straightforward: by avoiding a deceased individual, one can decrease the odds of encountering the threat that led to their demise.

Various species across phyla respond to diverse death cues. Eusocial species such as ants detect chemical cues on the carapaces of living colony members to distinguish them from dead conspecifics, which they subsequently remove from the colony in a process called ‘necrophoresis’^2^. *Drosophila* visually recognize other dead flies, particularly conspecifics, and can differentiate between animals that have died from hazardous sources (infection) as compared to flies that don’t “look dead” (flash frozen)^1^. Zebrafish exhibit panic behaviors in the presence of ‘Schreckstoff’ (fear substance), a mixture of multiple olfactants released from the injured skin of other fish^3–5^. Even at the scale of single cells, death signals are sufficient to induce ‘necrotaxis’, i.e., directed movement in response to “the agony” of a dead or dying cell^9^. Necrotaxis was originally observed in vertebrate immune cells, which necrotax towards (‘positive necrotaxis’) dying cells to perform phagocytosis^9^ and promote wound healing^10^. By contrast, multiple species of unicellular organisms perform ‘negative necrotaxis’ and avoid the lysed remains of cells^11,12^.

The nematode *Caenorhabditis elegans* is a powerful system with which to dissect the fundamental molecular biology underpinning both individual and social behaviors due to its highly invariant, stereotyped, and specialized nervous system^13^. Even with only 302 neurons, *C. elegans* is capable of complex decision-making when foraging in its environment to promote survival. *C. elegans* recognizes multiple cues in the remains of other dead worms, including olfactory cues sensed by the AWB neurons and “alarm pheromone” sensed by the ASI and ASK neurons^7,14^.

Here, we describe a novel gustatory necrotaxis cue, which we term Todstoff (‘death substance’), present in the remains of deceased *C. elegans* that is distinct from previously-identified worm pheromones and death cues. Our results demonstrate that Todstoff is sensed by GPCRs expressed in the ASH nociceptive neurons, which transduce signals to the AIB interneurons via the G-alpha protein ODR-3, TRPV channel orthologs OSM-9 and OCR-2, and synaptic glutamatergic signaling to induce avoidance. Todstoff co-precipitates with proteins and can be biochemically fractionated, enabling its isolation. Through this analysis, we identified Todstoff as a protein-associated small molecule. Taken together, our work illuminates a novel mode of post-mortem inter-animal chemical communication to promote death avoidance.

## Results

### *C. elegans* performs negative necrotaxis in response to the lysed remains of dead conspecifics

Motile organisms must integrate information from danger-associated and appetitive cues when making decisions in navigating their environment. In particular, cues associated with the death of conspecifics can be potent signals that drive avoidance even if a highly attractive target (such as a plentiful food source) is nearby. *C. elegans* robustly avoids the lysate of other *C. elegans* (Figure 1A)^7,8^. To quantify the strength of death signals over the attractive signals from food (*E. coli*), we designed a necrotaxis choice assay (Figure 1B) in which worms are placed onto an agar surface between two *E. coli* lawns with a line of either buffer or lysate separating them from each respective patch (Figure 1B; see Methods) and scored the number of worms that crossed one of the lines to reach a patch of food. Importantly, this assay: 1) measures the worms’ first choice (by paralyzing them with sodium azide when they reach an *E. coli* lawn) and does not permit them to change sides or learn to avoid lysate after prolonged exposure; and 2) presents lysate without the context of appetitive gustatory bacterial cues. We observed that worms robustly avoided the food patch on the side of the lysate barrier (Figure 1C). We also observed a strong avoidance of lysate made from *daf-22* ascaroside biosynthesis mutant *C. elegans* (Figure 1D), suggesting that the necrotaxis cue is distinct from known pheromone dispersal signals^15^. To avoid inadvertently complicating death avoidance from known pheromone avoidance pathways, we used *daf-22 C. elegans* lysate in all of our subsequent experiments.

**Figure 1.**
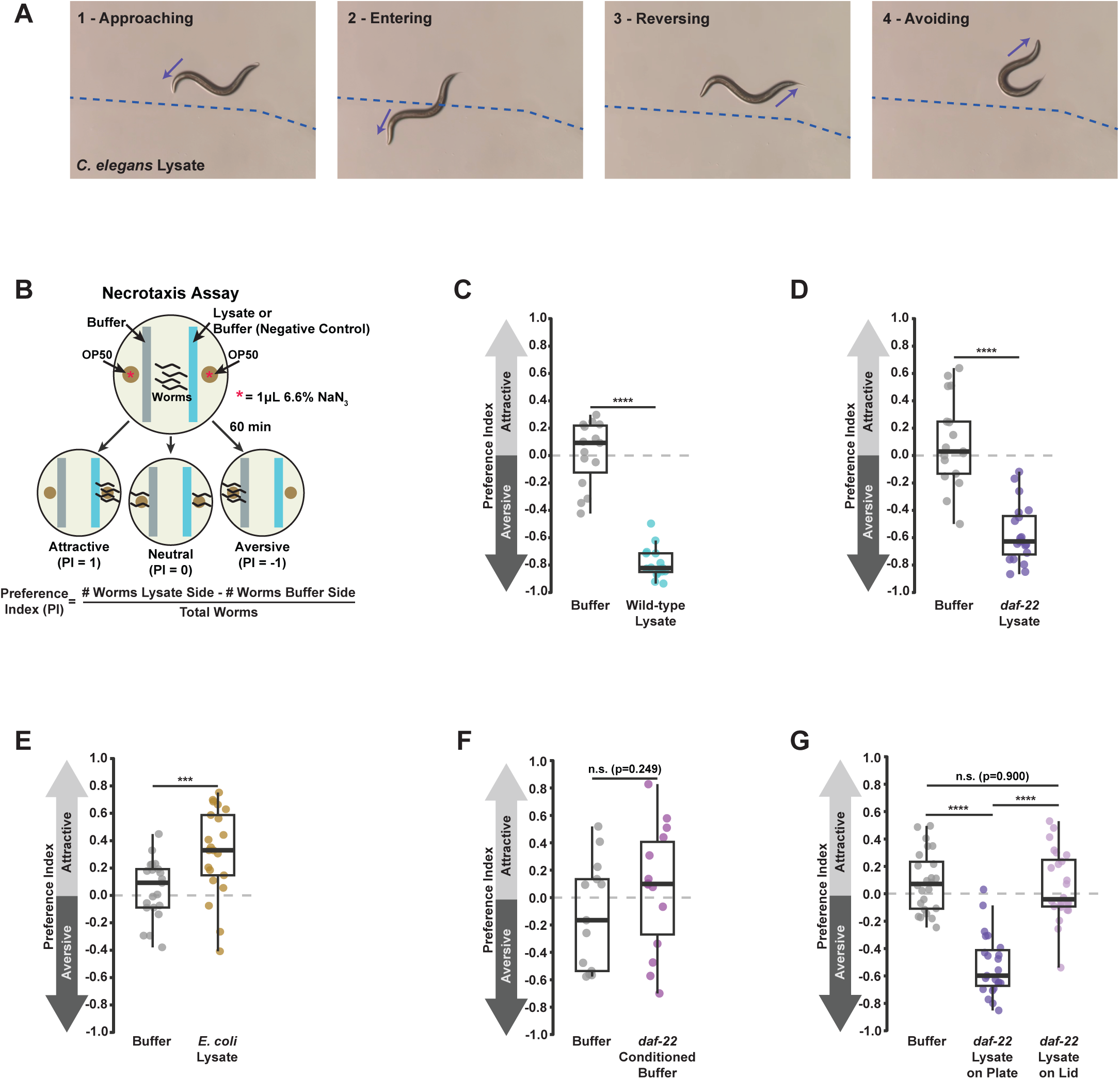
*C. elegans* performs negative necrotaxis to a gustatory cue in the remains of lysed conspecifics. A) Images taken from Supplemental Video 1 of a day 1 adult wild-type worm navigating on an agar plate upon which a pool of lysate from day 1 adult *daf-22 C. elegans* was placed and allowed to dry. The dashed line indicates the edge of the lysate, and arrows indicate the movement trajectory of the worm. B) Diagram of the necrotaxis assay. C) Necrotaxis assay of day 1 adult wild-type *C. elegans* responding to lysate from day 1 adult wild-type worms (3 combined experimental replicates). D) Necrotaxis assay of day 1 adult wild-type *C. elegans* responding to lysate from day 1 adult *daf-22* mutant *C. elegans* (3 combined experimental replicates). E) Necrotaxis assay of day 1 adult wild-type *C. elegans* responding to lysate from *E. coli* (3 combined experimental replicates). F) Necrotaxis assay of day 1 adult wild-type *C. elegans* responding to media conditioned with the secretions of day 1 adult *daf-22* hermaphrodites (2 combined experimental replicates). G) Necrotaxis assay of day 1 adult wild-type *C. elegans* responding to lysate from day 1 adult *daf-22* mutant *C. elegans* placed either on the agar or the lid of a plate (3 combined experimental replicates). P values in panels C-F were calculated by Student’s t test. P values in panel G were calculated by one-way ANOVA with Tukey’s HSD tests for multiple comparisons. * = p<0.05, ** = p<0.01, *** = p<0.001, **** = p<0.0001. In panels with box plots, the upper and lower hinges indicate the first and third quartiles, while the whiskers indicate the minima and maxima (interquartile range ±1.5*IQR), and horizontal lines indicate the median.

Worms did not avoid lysate from *E. coli*, indicating that the necrotaxis cue is not ubiquitously present in all cell lysates (Figure 1E). We also did not observe avoidance of buffer conditioned with living *daf-22* mutant *C. elegans* (Figure 1F), demonstrating that the aversive cue is not a secreted factor. Finally, we tested whether the aversive cue was olfactory^14^ by assessing whether worms would avoid lysate when it was placed on the lid of the petri plate, rather than on the agar. Worms avoided lysate placed on the agar, but not lysate placed on the lid (Figure 1G), indicating that the necrotaxis cue is likely gustatory rather than olfactory.

### Identification of ‘Todstoff’ - a novel necrotaxis cue

*C. elegans* avoids at least two distinct danger signals present in the bodies of other worms - an ASI/ASK sensed “alarm pheromone”^7^ and a multi-component AWB-sensed death odor^14^. In our necrotaxis assay, we observed robust avoidance from strains in which the ASI and ASK neurons were ablated by caspase expression (Figure 2A-B) as well as AWB-defective *ceh-37* mutants^16^ (Figure 2C). We similarly found that neither AMP nor histidine, two of the AWB-sensed death avoidance signals detected by *C. elegans*^8^, was sufficient to induce avoidance in our necrotaxis assay (Figure 2D-E) even at concentrations much higher than physiological levels. These data suggested to us that we had identified a novel, third necrotaxis signal in *C. elegans*.

**Figure 2.**
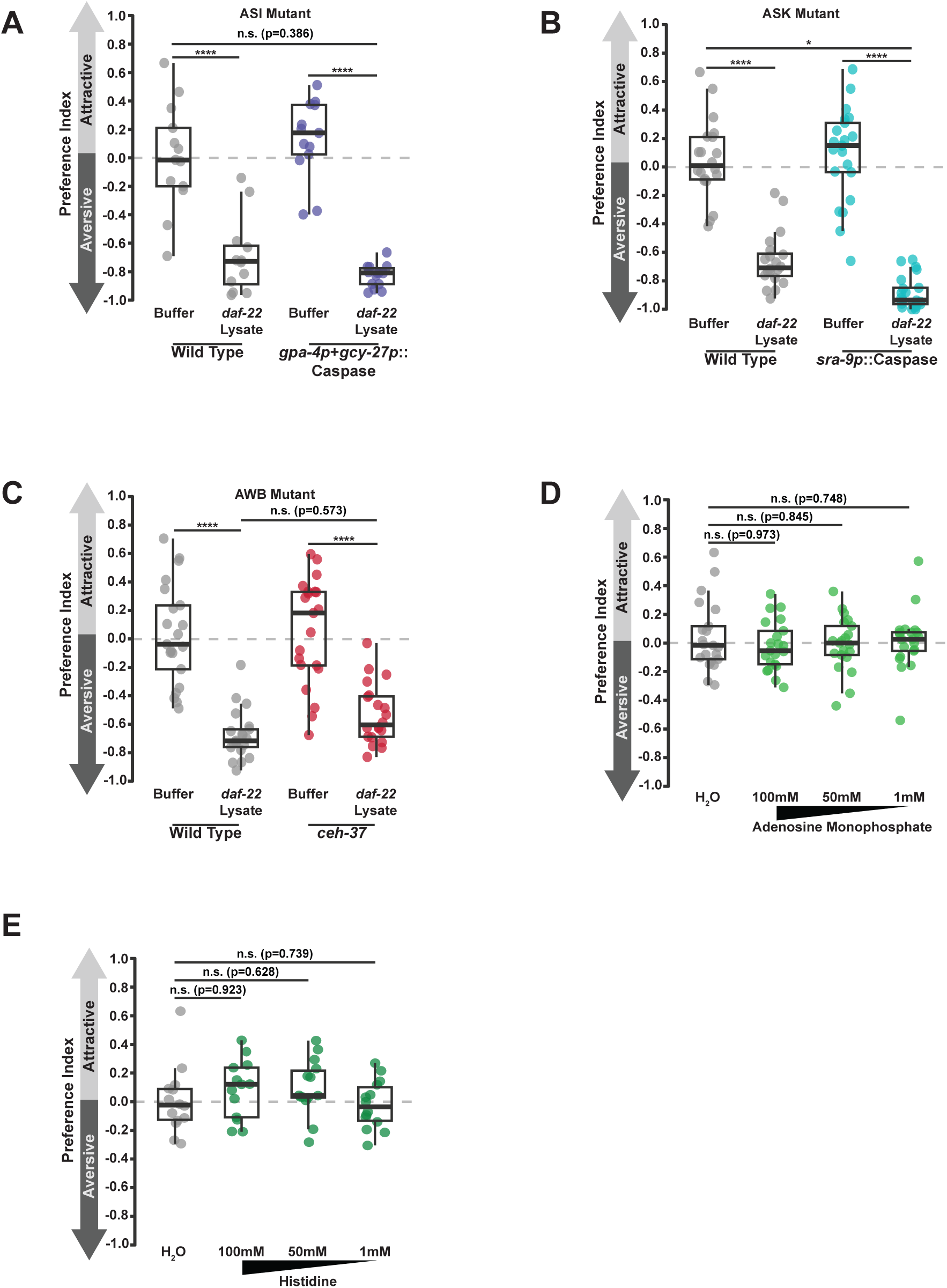
Identification of a novel necrotaxis cue. A-C) Necrotaxis assay of day 1 adult wild type, pASI::caspase (A), pASK::caspase (B), and *ceh-37* AWB mutant (C) *C. elegans* responding to lysate from day 1 adult daf-22 mutant *C. elegans*. D-E) Necrotaxis assay of day 1 adult wild type *C. elegans* responding to adenosine monophosphate (D) or histidine (E) in the necrotaxis assay. P values in panels A-C were calculated by two-way ANOVA with Tukey’s HSD tests for multiple comparisons. P values in panels D-E were calculated by one-way ANOVA with Tukey’s HSD tests for multiple comparisons. * = p<0.05, ** = p<0.01, *** = p<0.001, **** = p<0.0001. In panels with box plots, the upper and lower hinges indicate the first and third quartiles, while the whiskers indicate the minima and maxima (interquartile range ±1.5*IQR), and horizontal lines indicate the median.

In summary, we found that *C. elegans* performs negative necrotaxis in response to a non-ascaroside cue that is present within the bodies of other worms. Furthermore, this cue is distinct from other previously-described AWB- and ASI/ASK-sensed death-associated molecules. As *C. elegans* avoidance of this death cue is reminiscent of zebrafish fear responses to ‘Schreckstoff’ (‘fear substance’), a mixture of molecules released from the injured skin of other fish^3–5^, we therefore termed this new aversive worm cue ‘Todstoff’ (‘death substance’).

### Todstoff is sensed by ASH nociceptive neurons

As necrotaxis behavior commences rapidly upon encountering lysate (Figure 1A), we hypothesized that Todstoff is directly detected by sensory neurons. In concordance with this hypothesis, necrotaxis behavior towards lysate was reduced in *bbs-8* mutants and eliminated in *che-2* mutants, both of which disrupt the cilia required for sensory neuron function (Figure 3A)^17,18^. Therefore, Todstoff is most likely sensed by a ciliated sensory neuron.

**Figure 3.**
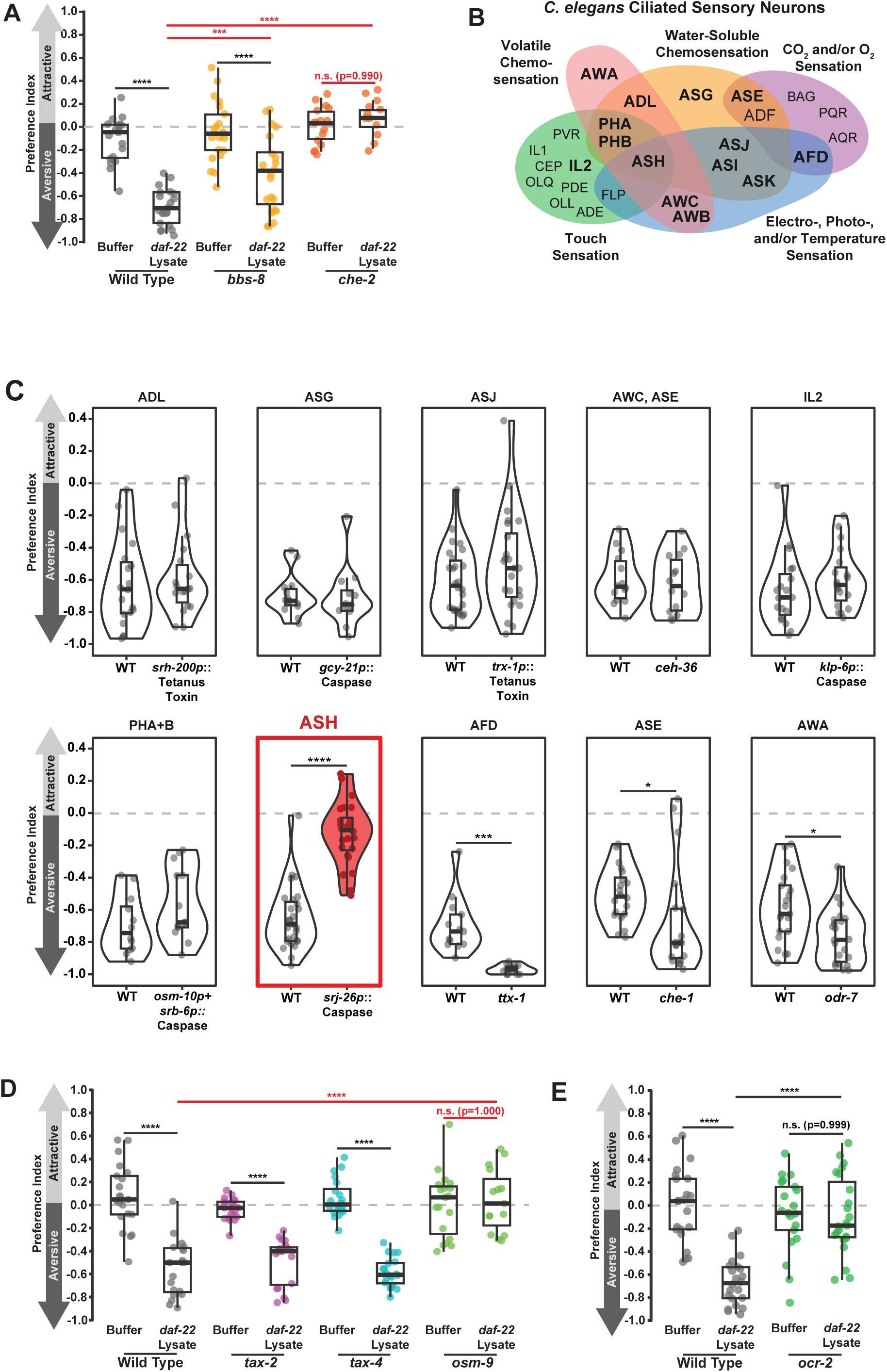
Todstoff is sensed by ASH nociceptive neurons. A) Necrotaxis assay of day 1 adult wild-type, *bbs-8*, and *che-2* mutant *C. elegans* responding to lysate from day 1 adult *daf-22* mutants (3 combined experimental replicates). B) Venn Diagram depicting the overall functions of ciliated neurons in *C. elegans*. Neuron functions were annotated from WormAtlas^19^. Bolded neurons were tested in this study. C) Necrotaxis assay candidate screen of mutants and genetic ablation lines disrupting specific *C. elegans* sensory neurons. Displayed are the preference indices for wild type and mutant worms on plates with PBS and Lysate. PBS vs PBS negative controls were also performed for each mutant (data not shown) and replicates were only used if the PBS controls between mutants were not statistically different by ANOVA with Tukey HSD tests for pairwise comparisons within replicates. Red violins and neuron names indicate mutants with significantly reduced avoidance relative to wild type (2-4 experimental replicates for each mutant). D) Necrotaxis assay of day 1 adult wild-type, *tax-2*, *tax-4*, and *osm-9* mutant *C. elegans* responding to lysate from day 1 adult *daf-22* mutants (3 combined experimental replicates). E) Necrotaxis assay of day 1 adult wild-type and *ocr-2* mutant *C. elegans* responding to lysate from day 1 adult *daf-22* mutants (2 combined experimental replicates). P values in panels A,D-E were calculated by one-way ANOVA with Tukey’s HSD tests for multiple comparisons. P values in panel C were calculated by Mann-Whitney U tests with Holm-Bonferroni correction. * = p<0.05, ** = p<0.01, *** = p<0.001, **** = p<0.0001. In panels with box plots, the upper and lower hinges indicate the first and third quartiles, while the whiskers indicate the minima and maxima (interquartile range ±1.5*IQR), and horizontal lines indicate the median.

*C. elegans* has distinct pairs of ciliated sensory neurons that perform specific functions (Figure 3B)^13,19^. To identify the sensory neuron(s) that detect Todstoff, we assessed the necrotaxis behavior of neuron-specific cell-fate mutants and genetic ablation lines for 10 different sensory neuron types (bolded in Figure 3B)^17,22–27^ in addition to the AWB, ASI, and ASK mutants that we previously tested (Figure 2A-C). Disruption of most sensory neurons had no effect on necrotactic avoidance of lysate (Figure 3C); however, ablation of the polymodal nociceptive ASH neurons significantly disrupted necrotaxis behavior, indicating that this neuron type is required for negative necrotaxis. The ASH neurons are the primary nociceptive sensory neurons in *C. elegans* that sense diverse repellant stimuli including touch, olfactants, water-soluble molecules, heavy metals, and osmolarity (Figure 3B)^13^. ASH chemosensation is not strongly impeded by defects in BBS-8-dependent intraflagellar transport, unlike other sensory neuron types^26,27^, providing rationale as to why *bbs-8* mutation only partially represses necrotaxis (Figure 3A).

By contrast, disruption of the function of the AFD (and more weakly, the ASE and AWA neurons) enhanced necrotactic avoidance (Figure 3C). AFD is known to reduce ASH sensitivity via the diffusion of cyclic nucleotides through gap junction channels^28^. Thus, the strong enhancing effect of AFD disruption on necrotaxis supports the model that ASH is the primary Todstoff-sensing neuron. AWA and ASE both sense food-related cues^13^, suggesting that perception of appetitive compounds may modulate Todstoff avoidance (Figure 2G-H).

Different classes of *C. elegans* neurons express distinct combinations of ion channels that are required for their function^13,29–31^. Avoidance of ASI/ASK and AWB-sensed death stimuli are dependent on the cyclic-nucleotide gated ion channels TAX-2 and TAX-4 and do not require the TRPV channel homolog OSM-9^7,8^. However, we found that mutation of *tax-2* or *tax-4* had no effect on necrotactic avoidance of Todstoff (Figure 3D). By contrast, mutation of the TRPV channels *osm-9* or *ocr-2* eliminated necrotaxis behavior (Figure 3D-E). As ASH expresses both *osm-9* and *ocr-2*, but not *tax-2* or *tax-4*^13^, these results further support the model that ASH neurons detect Todstoff, and that Todstoff-stimulated necrotaxis is distinct from other death avoidance behaviors.Together, these data demonstrate that Todstoff is sensed by the ASH neurons, but that necrotaxis behavior may be modulated by inputs from multiple sensory neurons.

### Molecular mechanism of necrotaxis signal transduction

Chemosensation in *C. elegans* is largely mediated by chemosensory G protein coupled receptors (csGPCRs)^13^. Our group’s single-nucleus sequencing of the adult *C. elegans* nervous system identified 147 csGPCRs that are expressed in ASH neurons; the majority of these GPCRs have no described function^32^. To efficiently identify the Todstoff csGPCR, we employed the recently developed worm GPCR mutant collection to test 88 strains carrying mutations for 610 csGPCRs^33^, including 99% (145/147) of ASH-expressed csGPCRs. Our screen yielded 5 candidate lines (Figure 4A), of which only one strain, CHS1233, exhibited defective necrotaxis in secondary testing (Figure 4B-C). While CHS1233 carries mutations for seven csGPCRs (*str-229*, *str-230, str-231, str-232, str-233, str-236,* C06B3.1, Figure 4A), only *str-230* is known to be expressed in the ASH^32,34^, highlighting this gene as the best candidate for the Todstoff receptor. *str-230* is also expressed in the ADL neurons; as we did not observe defective necrotaxis in an ADL-disrupted strain (Figure 3C), it is unlikely that ADL is also a primary sensor of Todstoff. However, ADL is a known secondary sensor of multiple nociceptive stimuli also detected by ASH^35–37^, suggesting that it may also serve as a secondary Todstoff sensor.

**Figure 4.**
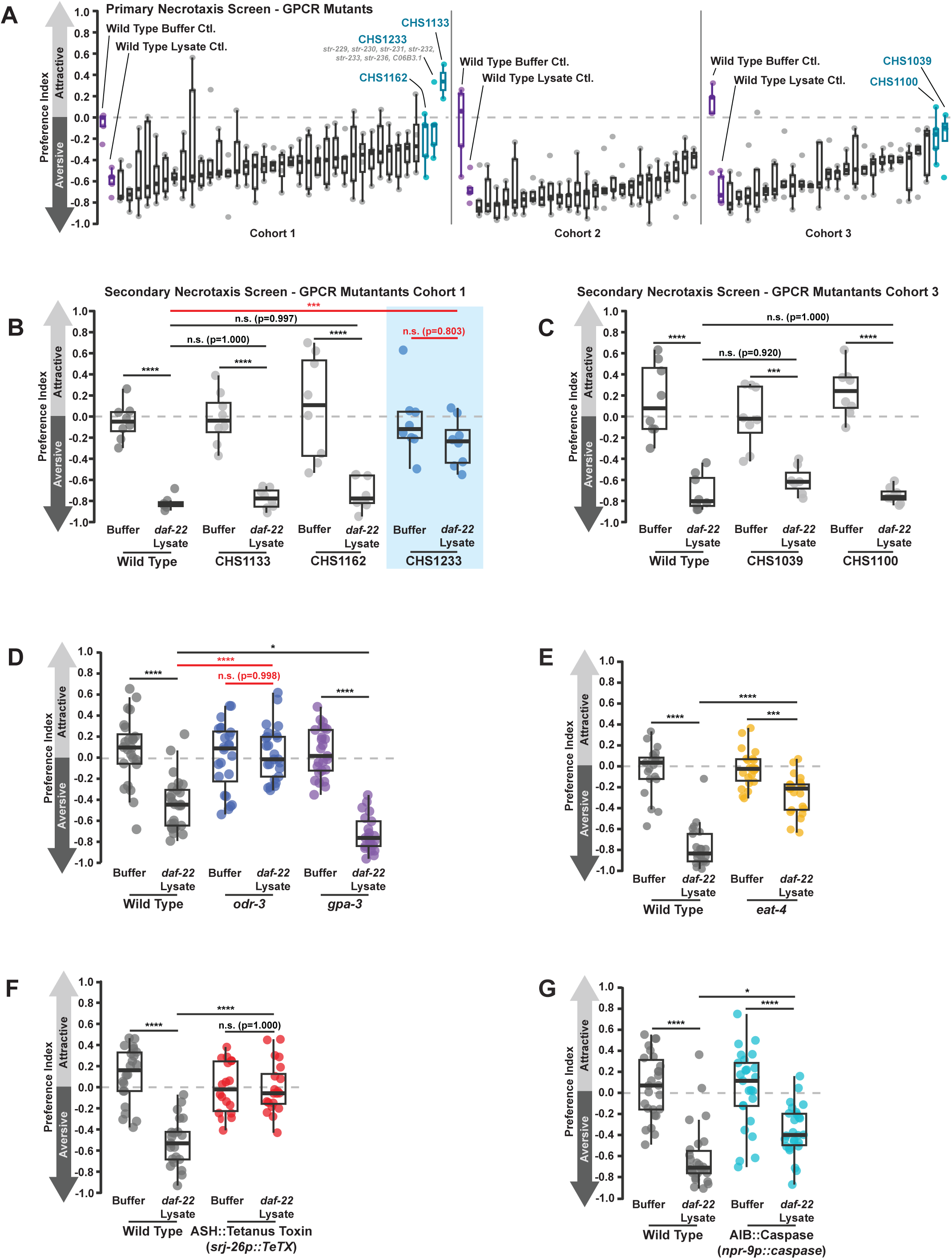
Molecular components of ASH signal transduction in necrotaxis. A) Primary screen of csGPCR mutants carrying mutations in GPCRs expressed in the ASH neurons ranked by median PI. 88 total mutants were tested across three experimental cohorts for necrotaxis to *daf-22* lysate in a 35mm plate assay (see Methods). Wild type controls and candidate strains with defective necrotaxis are labelec on the plot (aberrant necrotaxis mutants were defined as having a median PI >-0.2 and having at least one plate with a PI >0). B-C) Secondary screen of candidate GPCR mutant strains performed using a standard Necrotaxis barrier assay. Only CHS1233 exhibited defective necrotaxis (highlighted with a blue background). D) Necrotaxis assay of day 1 adult wild-type, *odr-3*, and *gpa-3* mutant *C. elegans* responding to lysate from day 1 adult *daf-22* mutants (2 combined experimental replicates). E) Necrotaxis assay of day 1 adult wild-type and *eat-4* mutant *C. elegans* responding to lysate from day 1 adult *daf-22* mutants (1 experimental replicate). F) Necrotaxis assay of day 1 adult wild-type and *ASH::Tetanus Toxin* expressing *C. elegans* responding to lysate from day 1 adult *daf-22* mutants (3 experimental replicates). G) Necrotaxis assay of day 1 adult wild type and *AIB::Caspase* expressing *C. elegans* responding to lysate form day 1 adult *daf-22* mutants (3 experimental replicates). In panels B-G, p values were calculated by two-way ANOVA with Tukey’s HSD tests for multiple comparisons. * = p<0.05, ** = p<0.01, *** = p<0.001, **** = p<0.0001. In panels with box plots, the upper and lower hinges indicate the first and third quartiles, while the whiskers indicate the minima and maxima (interquartile range ±1.5*IQR), and horizontal lines indicate the median.

GPCR signaling is mediated by the activity of G-proteins, and accordingly most ASH-mediated chemosensation requires the Gi/o-like proteins ODR-3 and/or GPA-3^38^. To determine which G protein(s) are required for Todstoff signal transduction, we examined the necrotaxis behavior of *odr-3* and *gpa-3* mutants (Figure 4D). We found that mutation of *odr-3*, but not *gpa-3*, was sufficient to prevent necrotaxis (Figure 4D). These data therefore support that Todstoff is sensed by a GPCR, which subsequently activates ODR-3 to initiate necrotaxis.

Sensory neurons can inform behavior both through chemical signaling at synapses and paracrine signals released from dense core vesicles^13^. ASH is glutamatergic, and the release of glutamate from ASH synapses is required for some, but not all, ASH-dependent avoidance behaviors^39^. To test whether synaptic transmission from ASH was required for necrotaxis, we examined the necrotaxis behavior of strains mutant for the vesicular glutamate transporter *eat-4* or expressing tetanus toxin specifically in the ASH, which prevents synaptic vesicle release (Figure 4E-F)^25,40,41^. Both ASH-expressed tetanus toxin and *eat-4* mutation were sufficient to disrupt Todstoff avoidance, indicating that glutamatergic synaptic communication with downstream interneurons is vital for necrotactic avoidance (Figure 4E-F).

ASH shares chemical synapses with multiple downstream interneurons. We found that ablation of the AIB interneurons, which function in the *C. elegans* ASH-stimulated reversal circuit^42^, partially repressed necrotaxis (Figure 4G). This phenotype suggests that the AIB interneurons contribute to necrotactic avoidance, but that they likely function in parallel with other interneurons to promote this behavior.

In summary, our data suggest that Todstoff sensation is a novel paradigm of death avoidance, and we have defined the neural pathway underlying its sensation, sensory transduction, and downstream neuron-neuron communication. Todstoff is sensed by the ASH neurons, which engage ODR-3-dependent GPCR signaling and transduce signals via the TRPV channels OSM-9 and OCR-2, culminating in glutamatergic synaptic signaling to AIB interneurons to promote necrotactic avoidance.

### Identification of Todstoff: Todstoff is not HPLA or EPA

We next set out to determine the molecular identity of Todstoff. *C. elegans* synthesizes biomolecules that can induce ASH-dependent avoidance^43,44^. Injured and infected *C. elegans* upregulate the synthesis of 4-hydroxyphenyllactic acid (HPLA), a tyrosine derivative that is sensed by the G protein-coupled receptor DCAR-1^44^*. dcar-1* is expressed in both the intestine and a subset of sensory neurons, including ASH, and is required for avoidance behaviors in response to water soluble cues analogous to HPLA^44,45^, raising the possibility that HPLA could serve as a necrotaxis cue. HPLA synthesis is not eliminated by genetic disruption of tyrosine metabolism pathways^44^, so to test whether the cue in Todstoff is HPLA, we assessed the necrotaxis behavior of *dcar-1* mutants (Figure 5A). We found that *dcar-1* mutants avoid lysate at levels indistinguishable from wild type, suggesting that DCAR-1 GPCR-mediated signaling and HPLA are not required for necrotaxis behavior.

**Figure 5.**
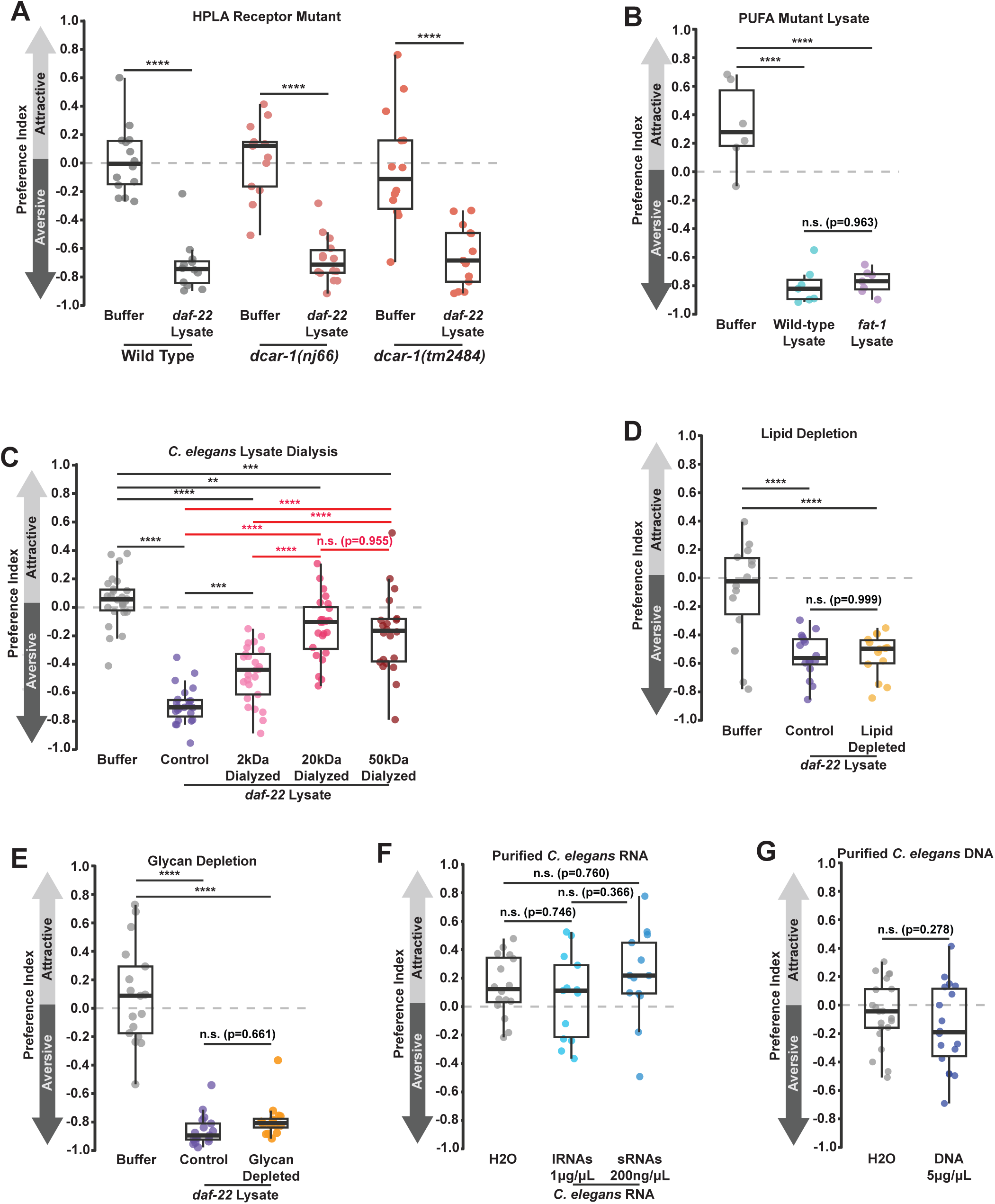
Candidate screen of biomolecules that may promote necrotaxis. A) Necrotaxis assay of day 1 adult wild-type and *dcar-1* mutant *C. elegans* responding to lysate from day 1 adult *daf-22* mutants. B) Necrotaxis assay of day 1 adult wild-type *C. elegans* responding to lysate from day 1 adult N2 or *fat-1* mutants (1 experimental replicate). C) Necrotaxis assay of day 1 adult wild-type *C. elegans* responding to lysate from day 1 adult *daf-22* mutants that has been dialyzed in membranes with molecular weight cutoffs of 2kDa, 20kDa, or 50kDa (3 combined experimental replicates). D) Necrotaxis assay of day 1 adult wild-type *C. elegans* responding to lysate from day 1 adult *daf-22* mutants that has been depleted of lipids with Cleanascite reagent (see methods, 2 combined experimental replicates). E) Necrotaxis assay of day 1 adult wild-type *C. elegans* responding to lysate from day 1 adult *daf-22* mutants that has been depleted of glycans with wheat germ agglutinin-coated beads (see methods, 2 combined experimental replicates). F) Necrotaxis assay of day 1 adult wild-type *C. elegans* responding to purified RNAs from day 1 adult N2 worms (2 combined experimental replicates). G) Necrotaxis assay of day 1 adult wild-type *C. elegans* responding to purified DNA from day 1 adult wild-type worms (2 combined experimental replicates). In panel A, p values were calculated by two-way ANOVA with Tukey’s HSD tests for multiple comparisons. In panels B-F, p values were calculated by Mann-Whitney U tests with Bonferroni-Holm correction for multiple comparisons. In panel G, the p value was calculated by Student’s t test. * = p<0.05, ** = p<0.01, *** = p<0.001, **** = p<0.0001. In panels with box plots, the upper and lower hinges indicate the first and third quartiles, while the whiskers indicate the minima and maxima (interquartile range ±1.5*IQR), and horizontal lines indicate the median.

*C. elegans* also synthesizes the polyunsaturated fatty acid eicosapentaenoic acid (EPA), which can induce *osm-9*-dependent avoidance behaviors^43^. We reasoned that this molecule may therefore induce necrotaxis. However, worms robustly avoided the lysate of *fat-1* worms, which do not synthesize EPA^43^, indicating that necrotaxis in response to Todstoff is distinct from EPA-induced avoidance (Figure 5B).

### Todstoff is smaller than 20 kDa

Our neuron mutant data on the sensation of Todstoff, in conjunction with our distinction of Todstoff from EPA and HPLA, suggested to us that Todstoff may be a previously unknown aversive molecule. To take an unbiased approach in characterizing Todstoff, we next approximated its size by dialyzing lysate from *daf-22* mutant worms, thereby preferentially depleting molecules smaller than a given molecular weight cutoff (MWCO) from the lysate (Figure 5C). Dialysis in a 20 kDa MWCO membrane significantly reduced aversion relative to both control lysate and lysate dialyzed in a 2 kDa MWCO membrane (Figure 5C). However, dialysis in a 50 kDa MWCO membrane did not further reduce aversion relative to the 20 kDa MWCO dialyzed lysate (Figure 5C). We also observed an intermediate reduction in lysate aversion following dialysis in a 2 kDa MWCO membrane relative to undialyzed control (Figure 5C); this result may indicate that Todstoff is near 2kDa in size, allowing inefficient diffusion through the membrane. In total, our dialysis data suggest that Todstoff is likely smaller in size than a 20 kDa protein.

### Todstoff is not composed of lipids, glycans, or nucleic acids

We next asked what class of biomolecule(s) composes Todstoff. Depletion of lipids from *daf-22 C. elegans* lysate using Cleanascite™ lipid depletion reagent did not prevent necrotaxis (Figure 5D, see Methods), nor did depletion of glycans (performed using wheat germ agglutinin (WGA) coated beads, see Methods) (Figure 5E). To test if Todstoff is composed of nucleic acids, we purified *C. elegans* RNA and DNA from lysate; however, worms did not avoid large RNAs, small RNAs, or DNA purified from lysate (Figure 5F-G), indicating that these molecules were not sufficient to induce necrotaxis.

### Todstoff precipitates with protein

We next used trichloroacetic acid (TCA) to precipitate protein^46^ from worm lysate and tested the re-solubilized pellet. We found that the resolubilized *C. elegans* lysate protein induced strong avoidance (Figure 6A). Importantly, TCA-precipitated protein from *E. coli* lysate did not exert the same effect (Figure 6A), indicating that *C. elegans* does not generally avoid precipitated and re-solubilized proteins. Avoidance of re-solubilized *C. elegans* protein is ASH-dependent (Figure 6B), indicating that we are following the same cue as in lysate. *C. elegans* protein precipitated with acetone was also aversive (Figure 6C), increasing our confidence that Todstoff precipitates with protein and that we were not observing an avoidance behavior that is artifactual to the precipitation method.

**Figure 6.**
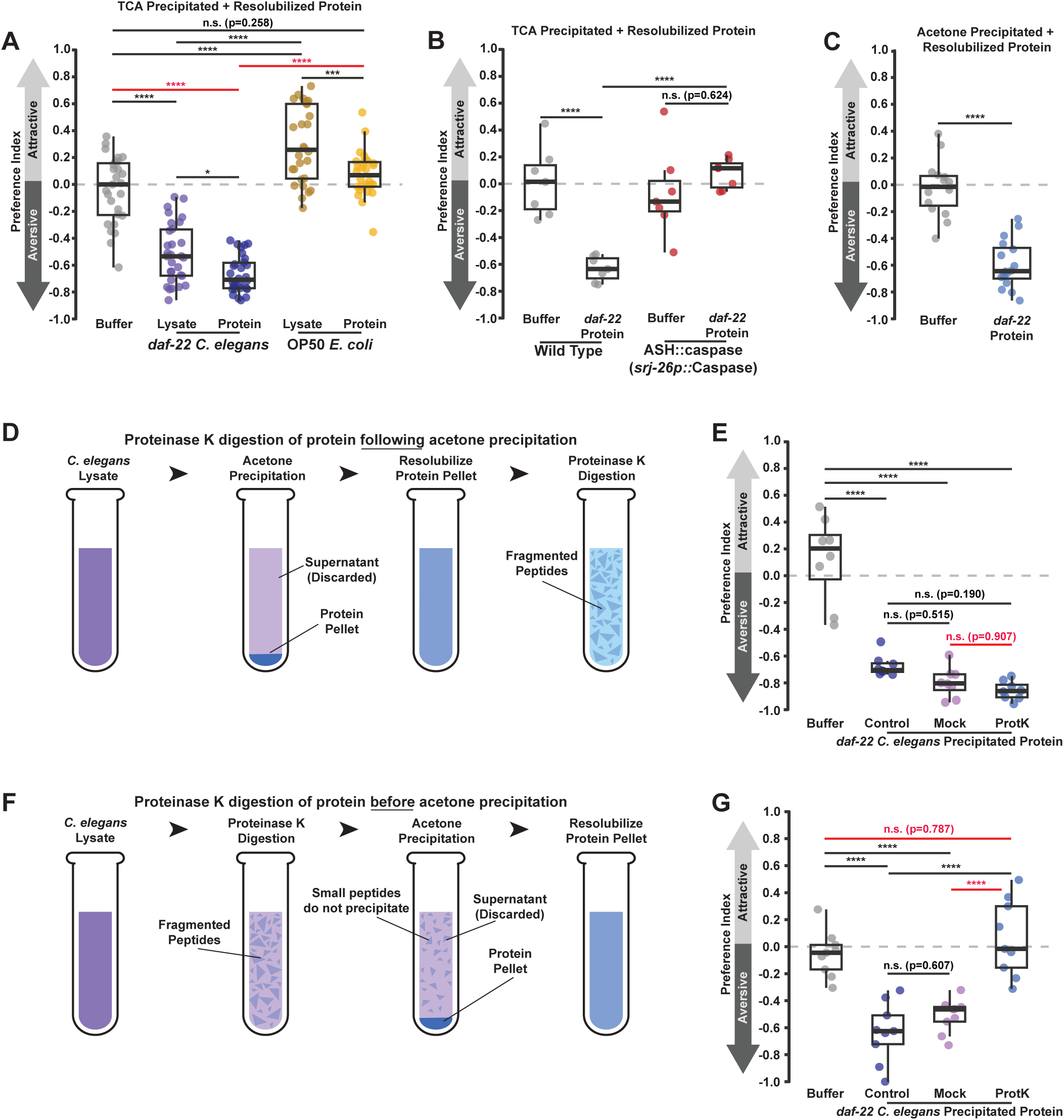
Todstoff precipitates with protein but is proteinase-resistant. A) Necrotaxis assay of day 1 adult wild-type *C. elegans* responding to protein from *daf-22 C. elegans* or OP50 lysate that was precipitated with trichloroacetic acid [TCA] and resolubilized (3 combined experimental replicates). B) Necrotaxis assay of day 1 adult wild-type and ASH::caspase expressing *C. elegans* responding to lysate or *daf-22 C. elegans* lysate protein precipitated with trichloroacetic acid [TCA] and resolubilized protein from *daf-22 C. elegans* worms or OP50 *E. coli* (1 experimental replicate). C) Necrotaxis assay of day 1 adult wild-type *C. elegans* responding to protein from *daf-22 C. elegans* lysate that was precipitated with acetone and resolubilized (3 combined experimental replicates). D) Cartoon depicting protocol for ProtK digestion of precipitated and resolubilized *C. elegans* protein. E) Necrotaxis assay of day 1 adult wild-type *C. elegans* responding to protein from *daf-22 C. elegans* or OP50 lysate that was first precipitated with acetone and resolubilized and then digested with Proteinase K (1 experimental replicate displayed of 3 total replicates). F) Cartoon depicting protocol of ProtK digestion of *C. elegans* lysate before protein precipitation and resolubilization. G) Necrotaxis assay of day 1 adult wild-type *C. elegans* responding to protein from *daf-22 C. elegans* or OP50 lysate that was first digested with Proteinase K and then precipitated with acetone and resolubilized (1 experimental replicate displayed of 3 total replicates). In panels A-B, p values were calculated by ANOVA with Tukey’s HSD tests for pairwise comparisons. In panel C, the p value was calculated by Student’s t test. In panels D-E, p values were calculated by 2 way ANOVA with Tukey’s HSD tests for pairwise comparisons. * = p<0.05, ** = p<0.01, *** = p<0.001, **** = p<0.0001. In panels with box plots, the upper and lower hinges indicate the first and third quartiles, while the whiskers indicate the minima and maxima (interquartile range ±1.5*IQR), and horizontal lines indicate the median.

We reasoned that if Todstoff is a protein, then degradation of proteins in lysate with Proteinase K (ProtK) should deplete the aversive cue (Figure 6D). However, we found that ProtK treatment of precipitated and resuspended *C. elegans* proteins had no effect on necrotaxis behavior (Figure 6E). This result suggested that Todstoff may be: 1) composed of a small peptide or domain that is not efficiently eliminated by proteinase K; 2) a proteinase-resistant protein; or 3) a non-protein molecule that co-precipitates with proteins.

Small peptides are inefficiently precipitated by most protein precipitation protocols^47^. We reasoned that if Todstoff is a small peptide or non-protein molecule that depends upon large proteins aggregating during precipitation to be retained, then ProtK treatment before precipitation should impede Todstoff precipitation (Figure 6F). Indeed, we observed that ProtK treatment of *C. elegans* lysate before acetone precipitation was sufficient to prevent Todstoff precipitation, as the resulting re-solubilized protein was not aversive (Figure 6G). Thus, our data suggest that although Todstoff co-precipitates with total protein in lysate, it is likely relatively soluble in conditions that precipitate proteins and requires association with or the aggregation of larger proteins to be retained in the precipitate.

### Fractionation of *C. elegans* protein isolates Todstoff

To separate Todstoff from other molecules present in precipitated and re-solubilized protein from worm lysate, we size-fractionated resolubilized worm protein via fast protein liquid chromatography (FPLC) and tested fractions encompassing molecules smaller than ∼17 kDa (in line with our determination of Todstoff’s approximate size by dialysis) for aversion. Of the 36 fractions tested, a contiguous subset of fractions eluting below the 1.35 kDa standard induced avoidance (Figure 7A), suggesting that those fractions contain Todstoff.

**Figure 7.**
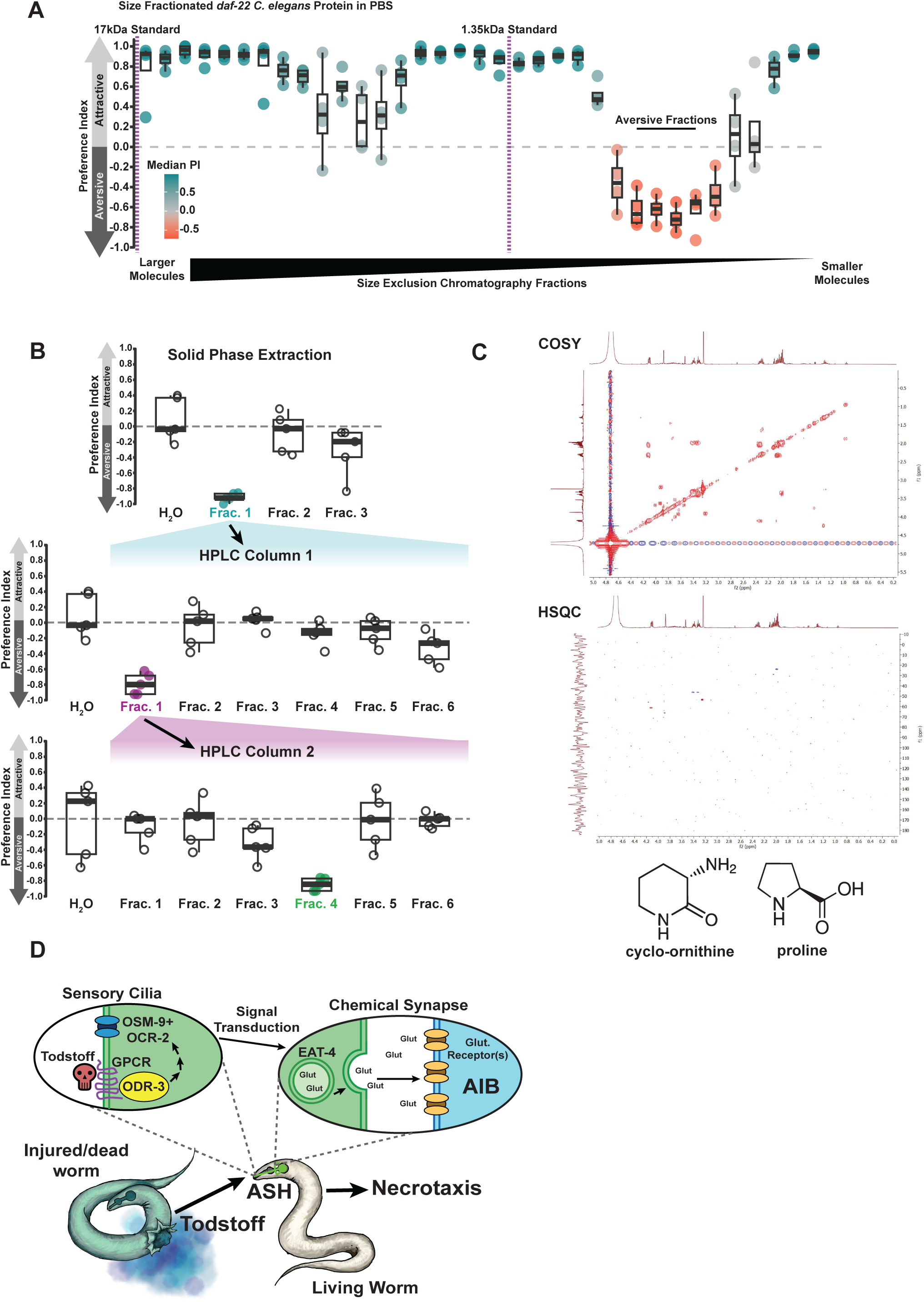
Biochemical isolation and fractionation of Todstoff. A) Necrotaxis assay of day 1 adult wild-type *C. elegans* responding to protein from *daf-22 C. elegans* that was fractionated by size-exclusion chromatography. The color of the individual datapoints represents the median preference index (Median PI) of that fraction. Inset panel displays UV280 absorbance of the fractions that induced avoidance. B) Pipeline by which precipitated and resolubilized *C. elegans* protein, which contains Todstoff, was processed to a minimal subset of molecules. Precipitated and resolubilized *C. elegans* protein was subjected to solid phase extraction followed by two rounds of reverse and normal phase HPLC. C) Preliminary correlation spectroscopy (COSY) and heteronuclear single quantum coherence (HSQC) NMR of aversive HPLC fractions reveals signal consistent with cyclo-ornithine or proline. D) Model of *C. elegans* Todstoff-induced necrotaxis. P values in panel C were calculated by one-way ANOVA with Tukey’s HSD tests for multiple comparisons. Model of *C. elegans* Todstoff-induced necrotaxis. * = p<0.05, ** = p<0.01, *** = p<0.001, **** = p<0.0001. In panels with box plots, the upper and lower images indicate the first and third quartiles, while the whiskers indicate the minima and maxima (interquartile range ±1.5*IQR), and horizontal lines indicate the median.

The very low approximate molecular weight of Todstoff suggested to us that it may not be a peptide, but rather a small molecule that co-precipitates with proteins. We therefore optimized an activity-guided biochemical fractionation pipeline to separate small molecules in our sample (Figure 7B). We first performed solid phase extraction (Figure 7B) and identified the aversive fraction using the necrotaxis assay. To further separate Todstoff from molecules that eluted with it, we performed reverse phase high-performance liquid chromatography (HPLC) on eluate from each of the UV-absorbing FPLC peaks (Figure 7B). We subjected this active fraction to two rounds of reverse- and normal-phase HPLC, again testing fractions using the necrotaxis assay to follow the active chemicals (Figure 7B). Using this approach, we were able to separate the compounds that co-precipitate with proteins to a minimal fraction. Our preliminary nuclear magnetic resonance spectroscopy analysis (NMR) revealed that this fraction contains multiple amino acids and amino acid derivatives, such as a signal that could be produced by cyclo-ornithine or proline (Figure 7C). Together, our data suggest that Todstoff is a small molecule (<1.35 kDa) that precipitates with proteins from lysed *C. elegans*, and causes potent avoidance by neighboring *C. elegans* via detection by the ASH nociceptive neuron, which uses a GPCR pathway and glutamatergic signaling to the AIB neuron.

## Discussion

Many species across phyla respond to death-specific stimuli, with many organisms sensing distinct cues to identify dead conspecifics. Here, we demonstrate that *C. elegans* avoids ‘Todstoff’, a novel necrotaxis cue present in the remains of other *C. elegans*. From our biochemical experiments, we provide evidence that Todstoff is a small molecule that precipitates with proteins (Figure 6A,7A). We find that the ASH polymodal nociceptors sense Todstoff through G protein coupled receptor signaling, activating the Gi/o-like protein ODR-3, which transduces this signal via the activation of the TRPV channels OSM-9 and OCR-2 to facilitate glutamatergic synaptic transmission to the AIB interneurons and avoidance behaviors (Figure 7D). Our work, in conjunction with studies by Hernandez-Lima et al. 2025 and Zhou et al. 2017, demonstrates that *C. elegans* perceives at least three modalities of death signaling: 1) ASH-sensed ‘Todstoff’; 2) ASI/ASK sensed “alarm pheromone”^7^; and, 3) AWB-sensed cues including AMP and Histidine^8^. This diversity in death cues may enable *C. elegans* to avoid hazardous contexts regardless of the modes of a conspecific’s death, thereby ensuring the living worm’s ability to avoid danger.

Nematode death signals, including Todstoff, may serve roles in not only intra-species communication but also inter-species conflict. Our work defining the Todstoff avoidance pathway will also enable us to survey its conservation across nematodes (and in non-nematode animals) to assess whether Todstoff or similar signals may serve as not only a *C. elegans* specific cue but as a danger signal more broadly across species. Taken together, we have defined a new modality of *C. elegans* alarm signaling paradigm by which animals perceive and avoid the remains of dead conspecifics.

## Methods

### *C. elegans* Strains and Culture Conditions

All *C. elegans* strains were maintained at 20°C under standard conditions on high growth medium (HG) agar plates [3g/L NaCl, 20g/L Bacto-peptone, 30g/L Bacto-agar, 4mL/L cholesterol (5 mg/mL in ethanol), 1 mL/L 1M CaCl_2_, 1 mL/L MgSO_4_, and 25mL/L 1M KPO_4_ pH6.0] seeded with OP50 *Escherichia coli* bacteria. Worm populations were synchronized by bleaching in alkaline-bleach solution [250mM KOH, 12% hypochlorite bleach] followed by two washes with M9 buffer [6 g/L Na_2_HPO_4_, 3g/L KH_2_PO_4_, 5 g/L NaCl, 1 mL/L 1M MgSO_4_ in ddH2O].

Strains used in this study include:

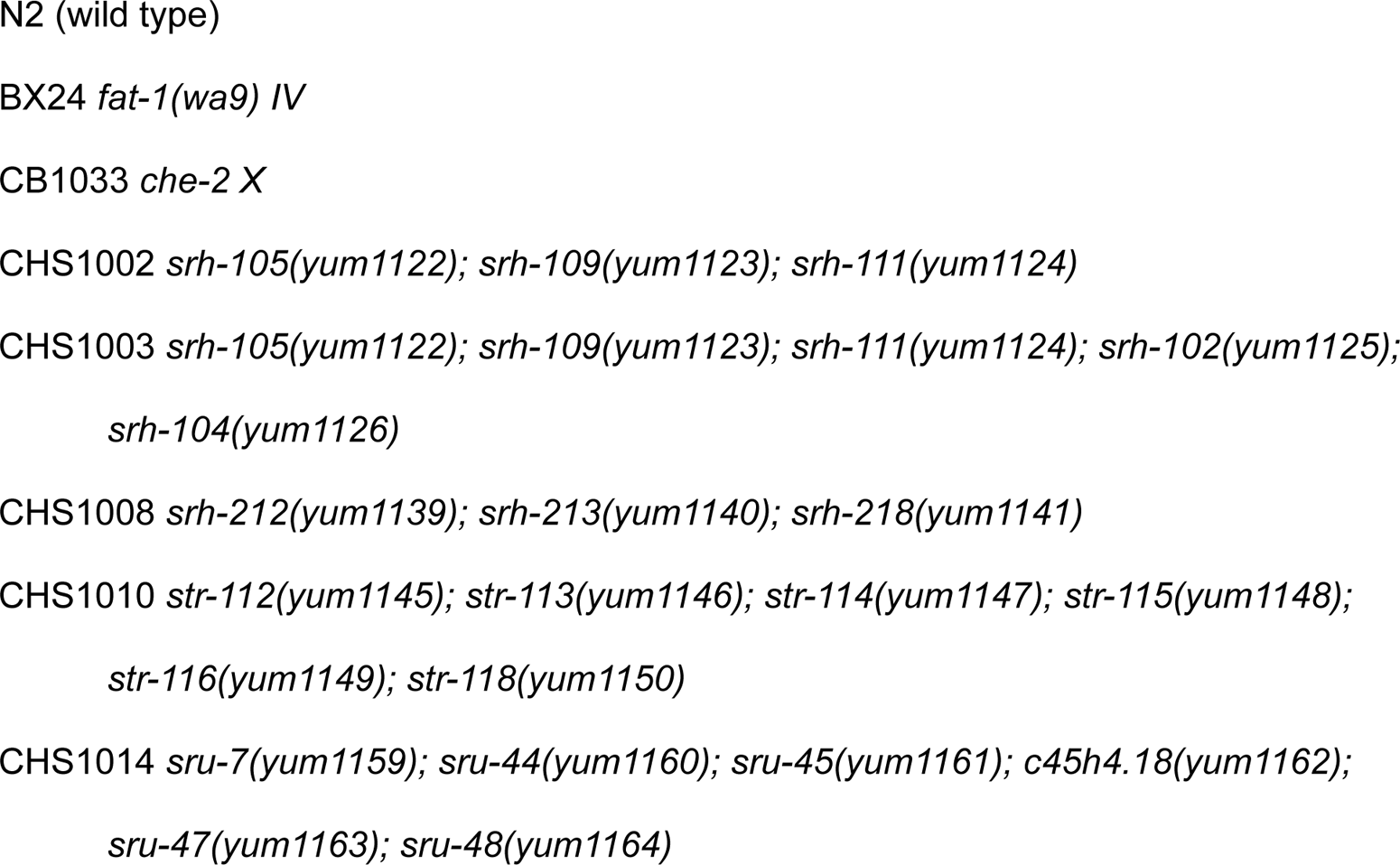
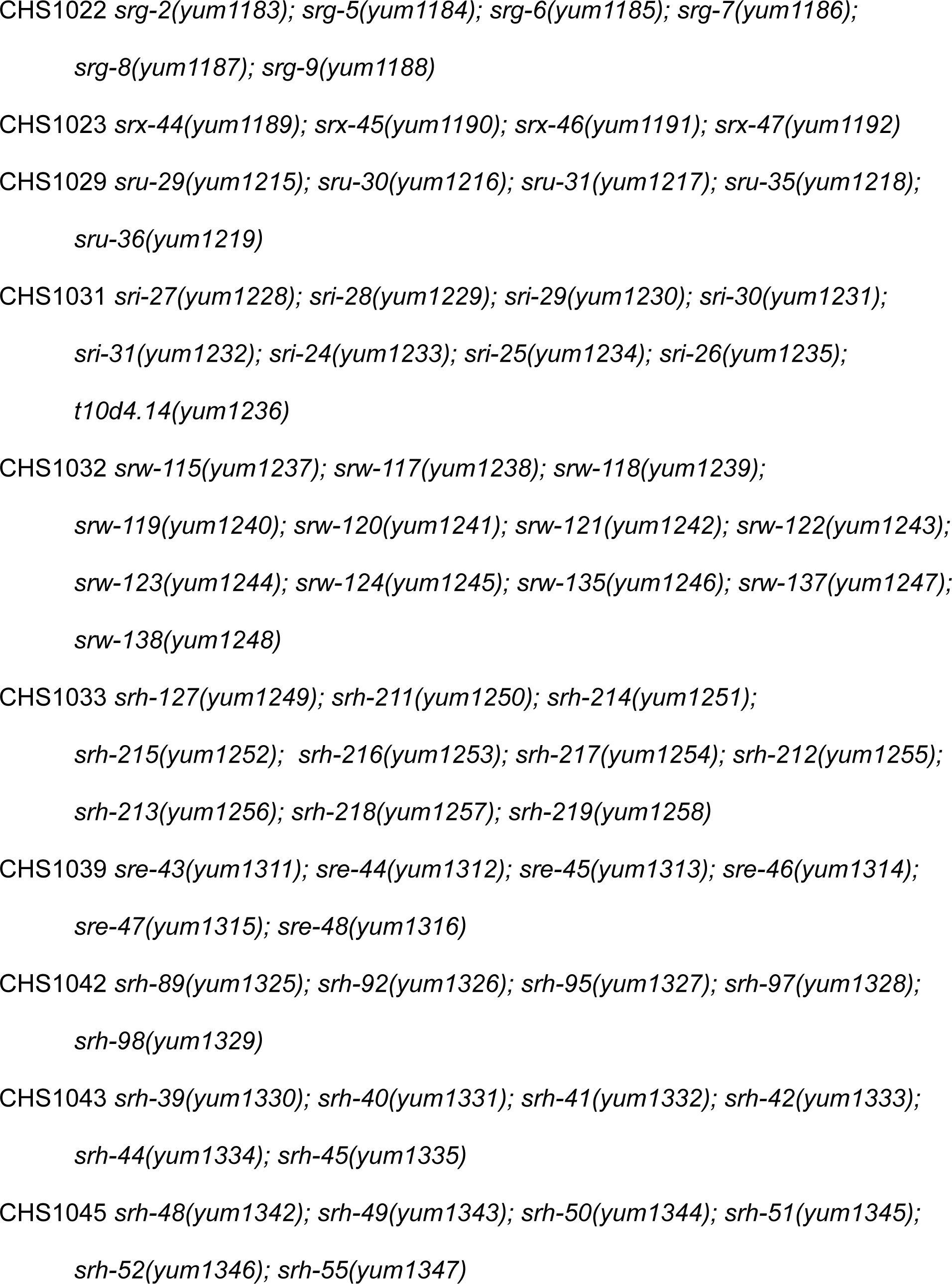
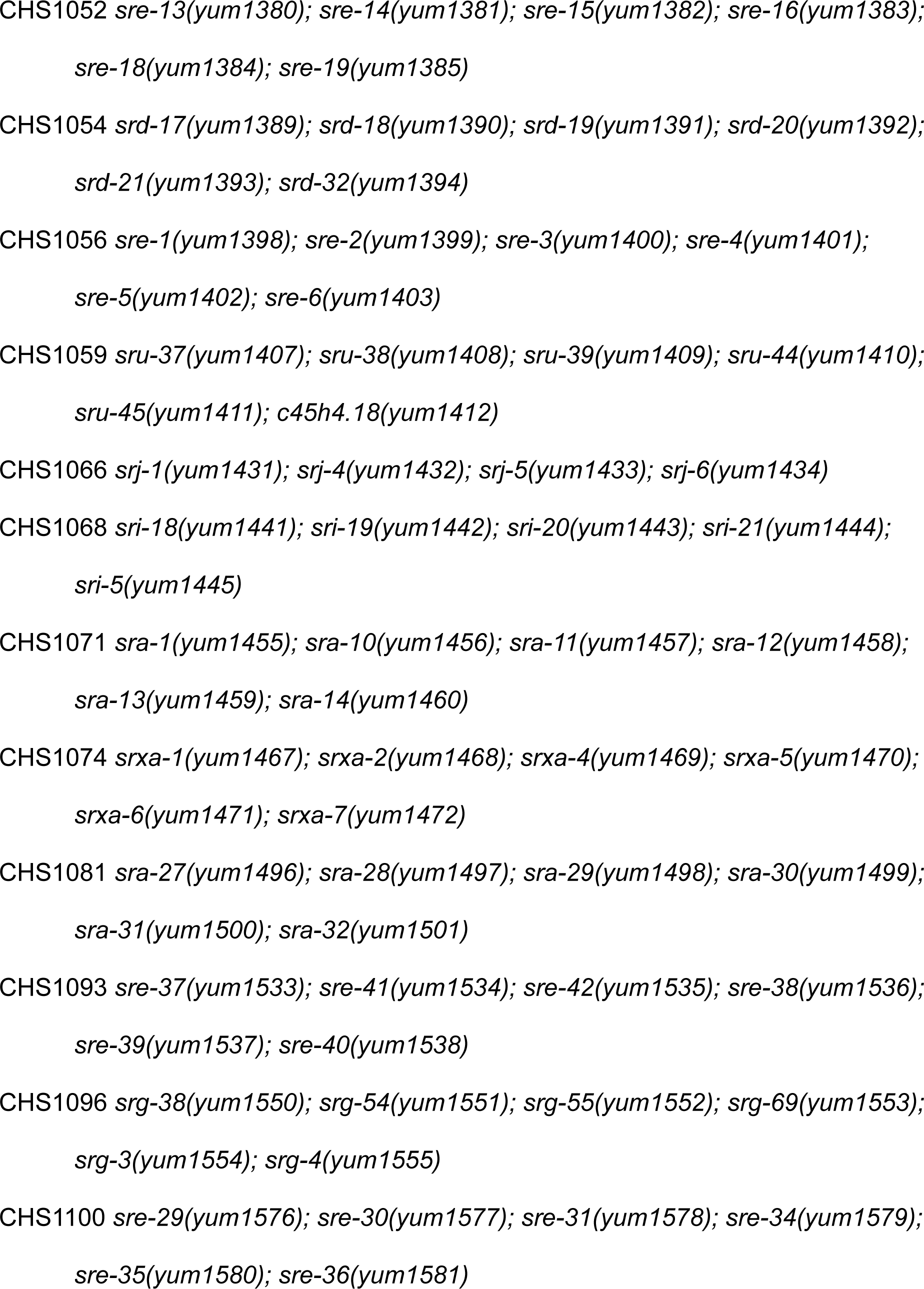
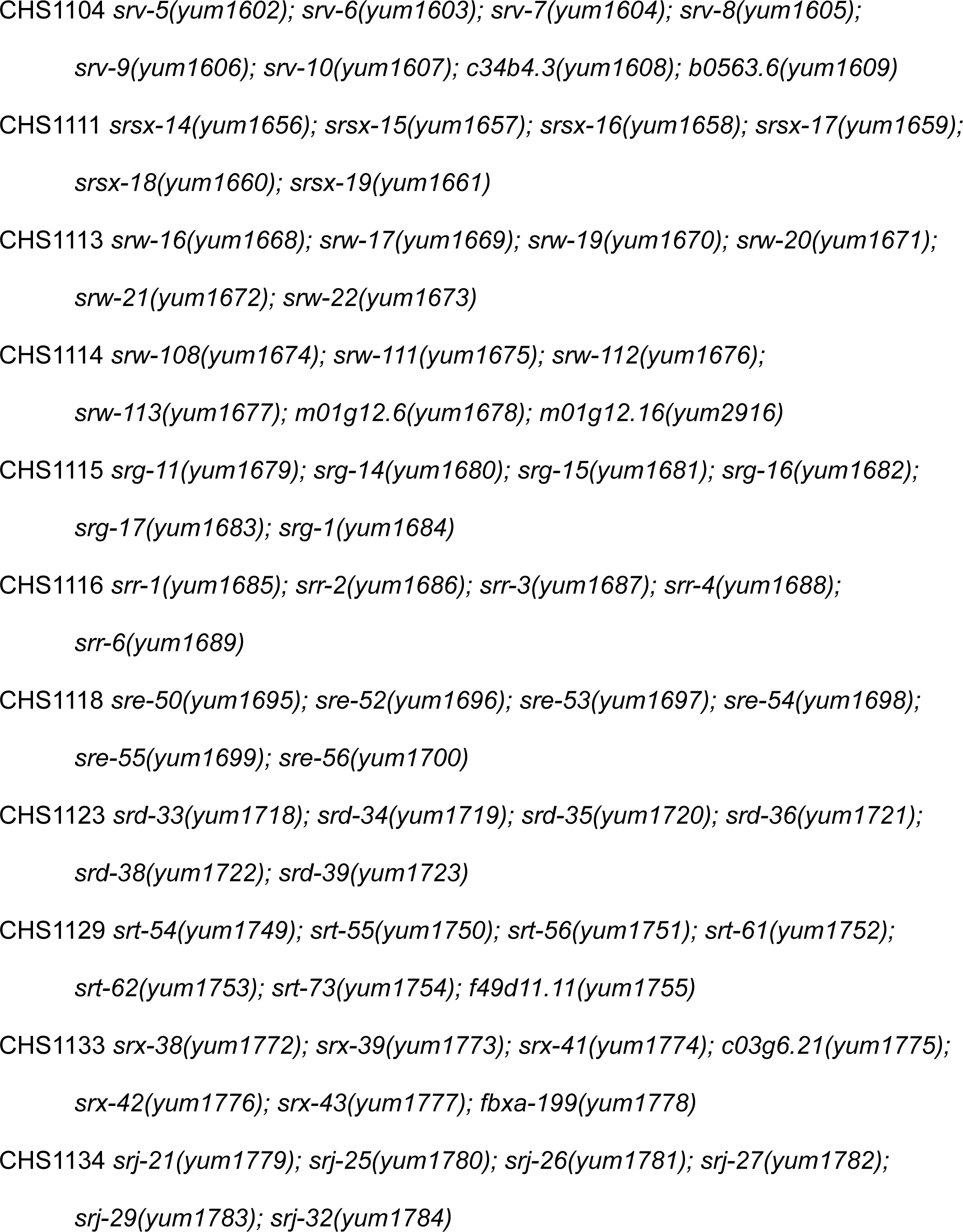
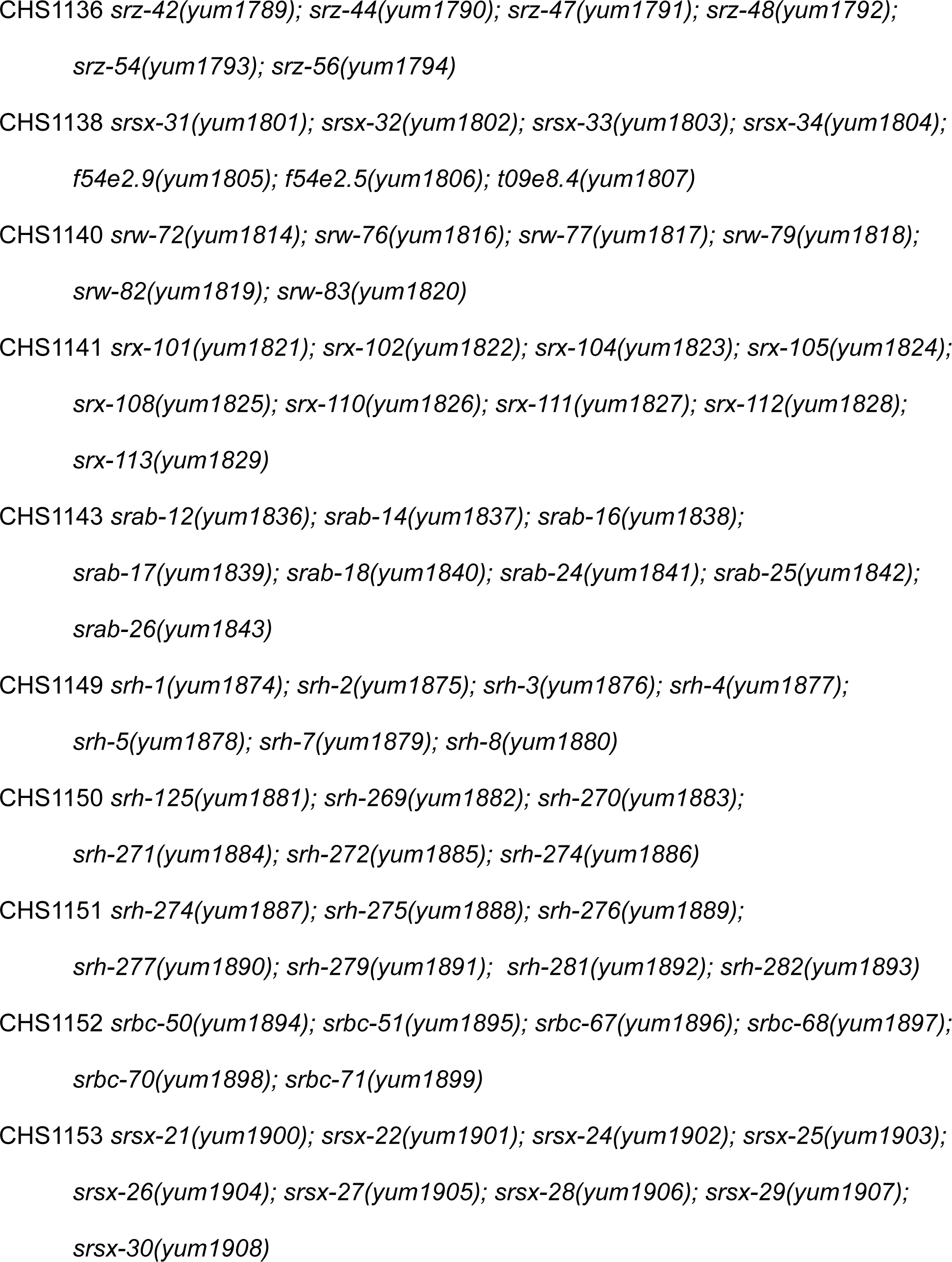
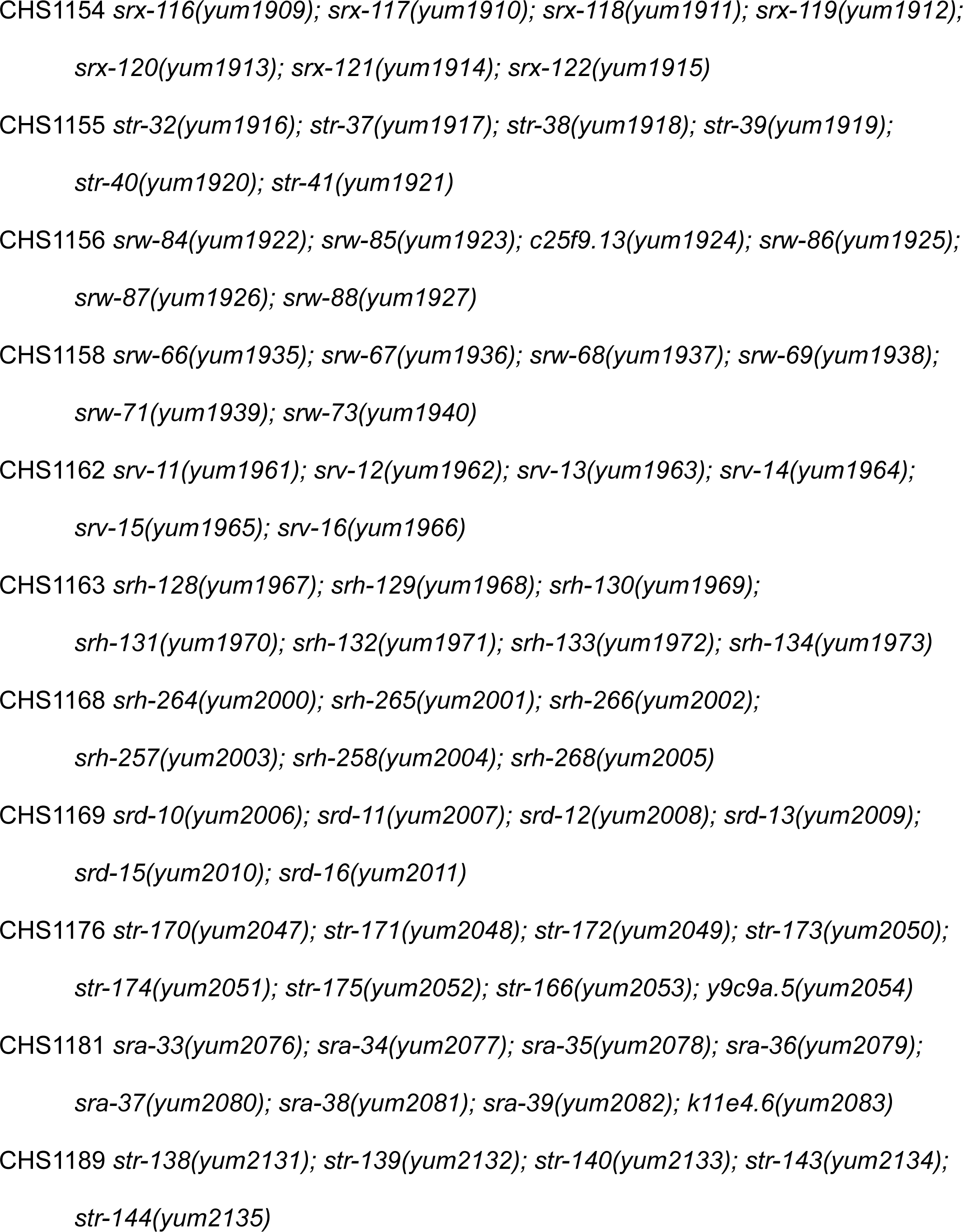
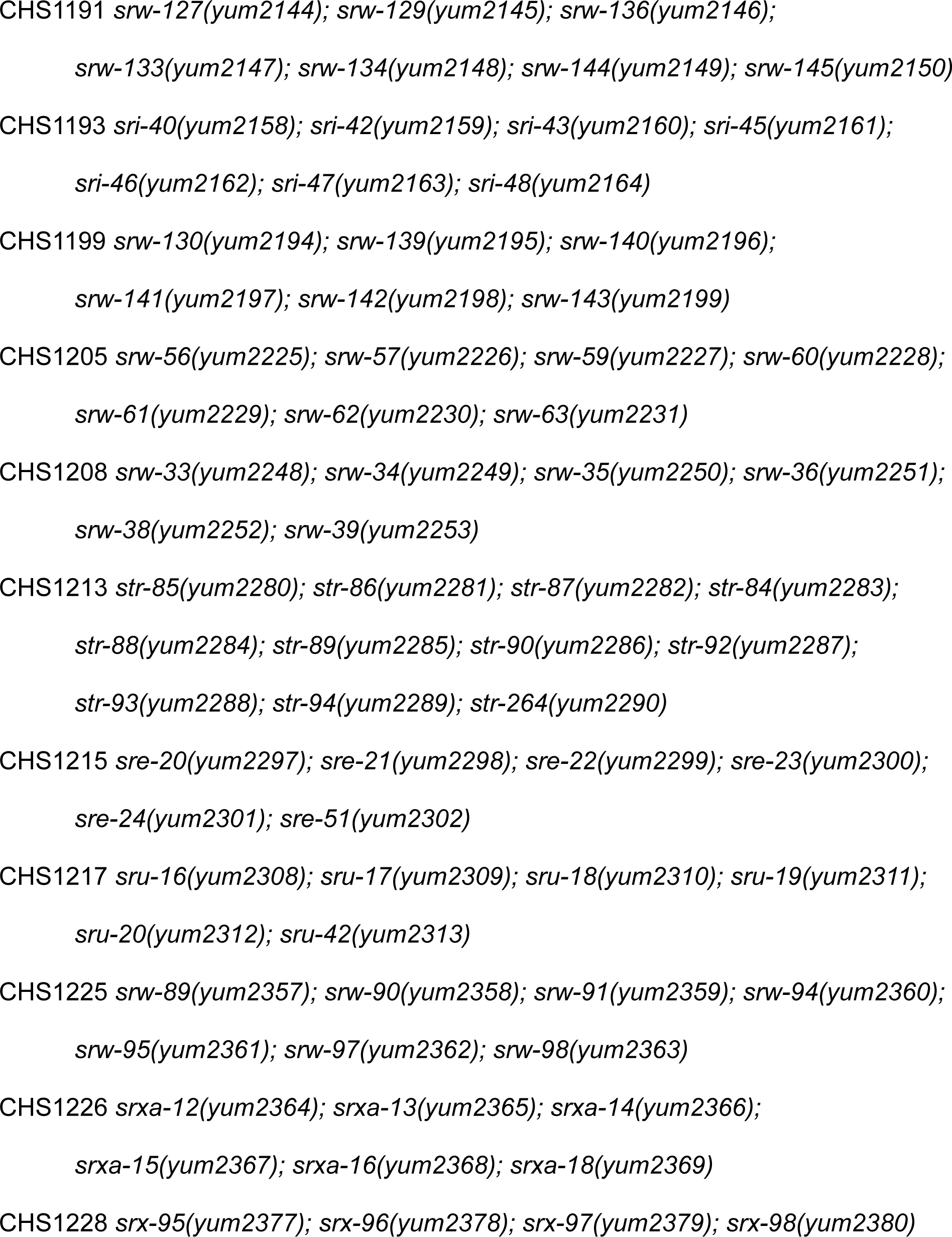
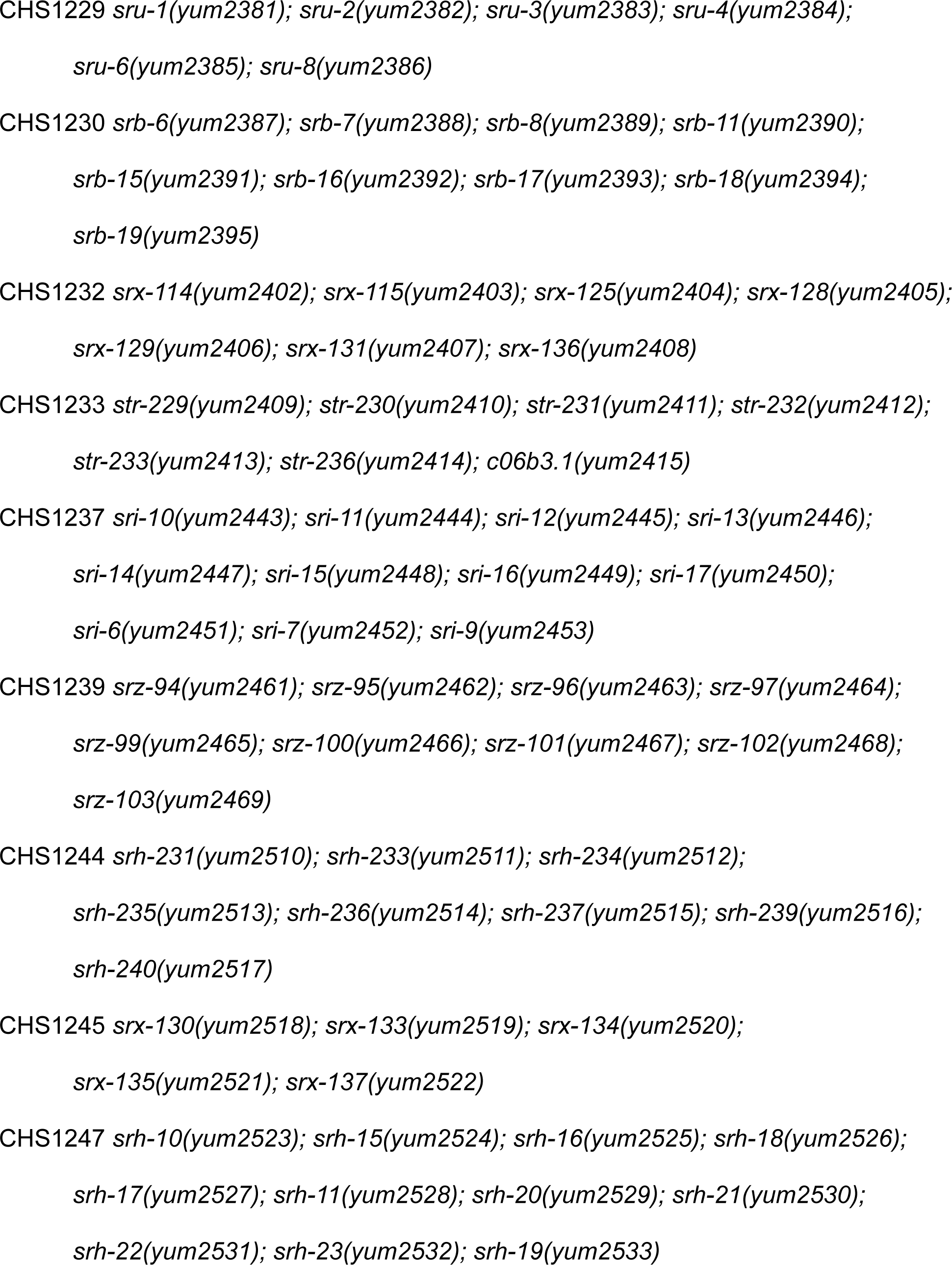
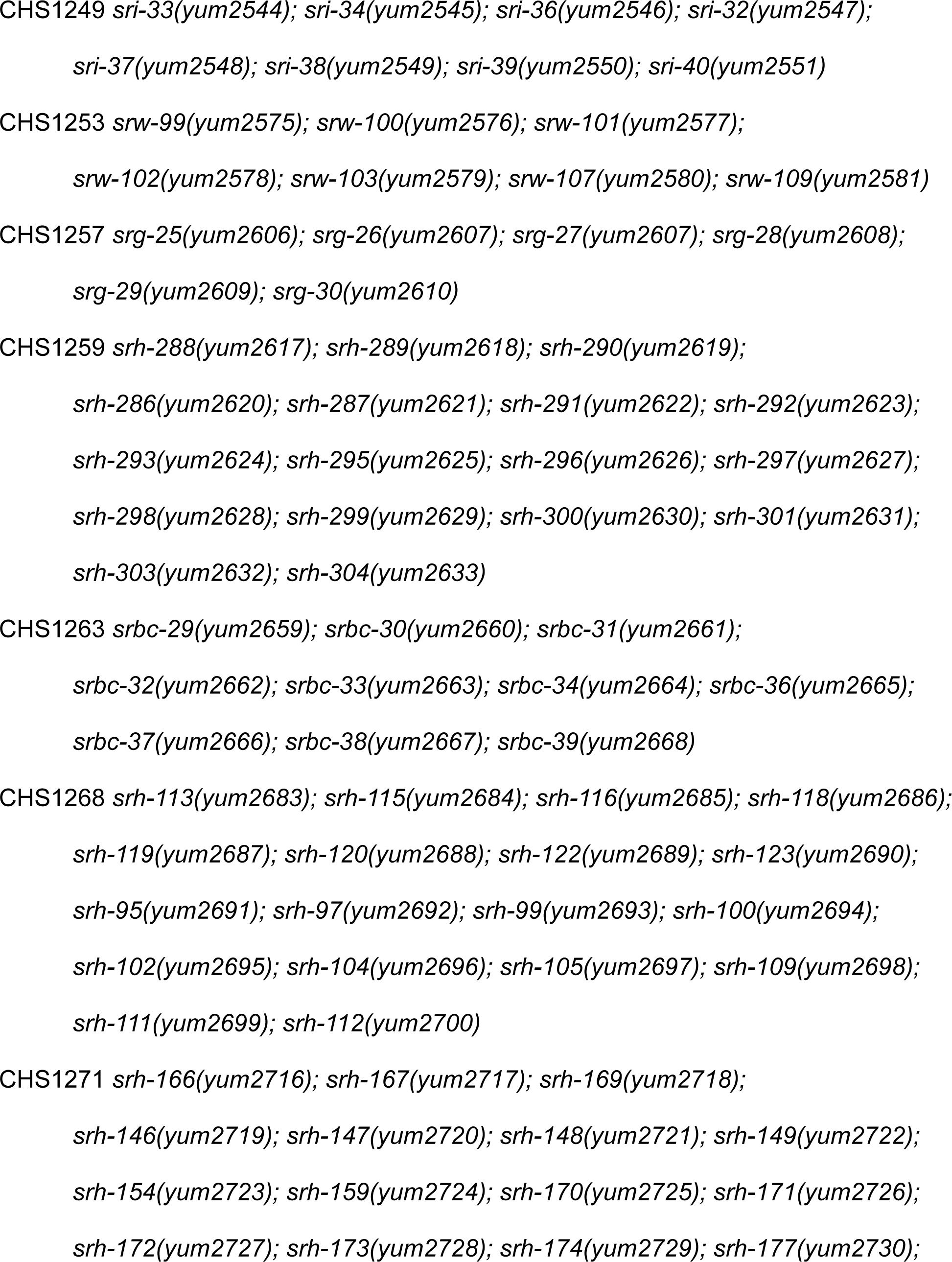
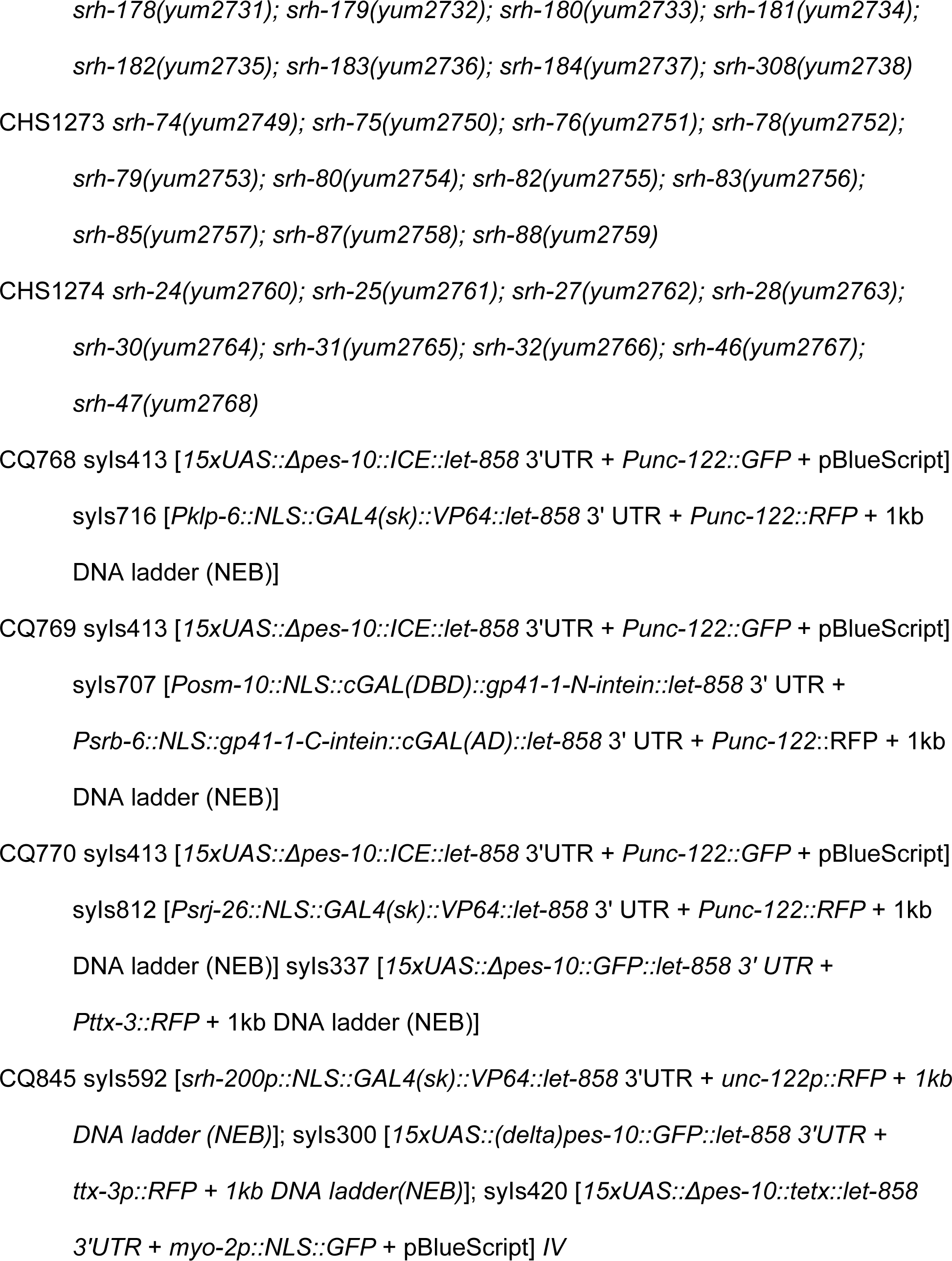
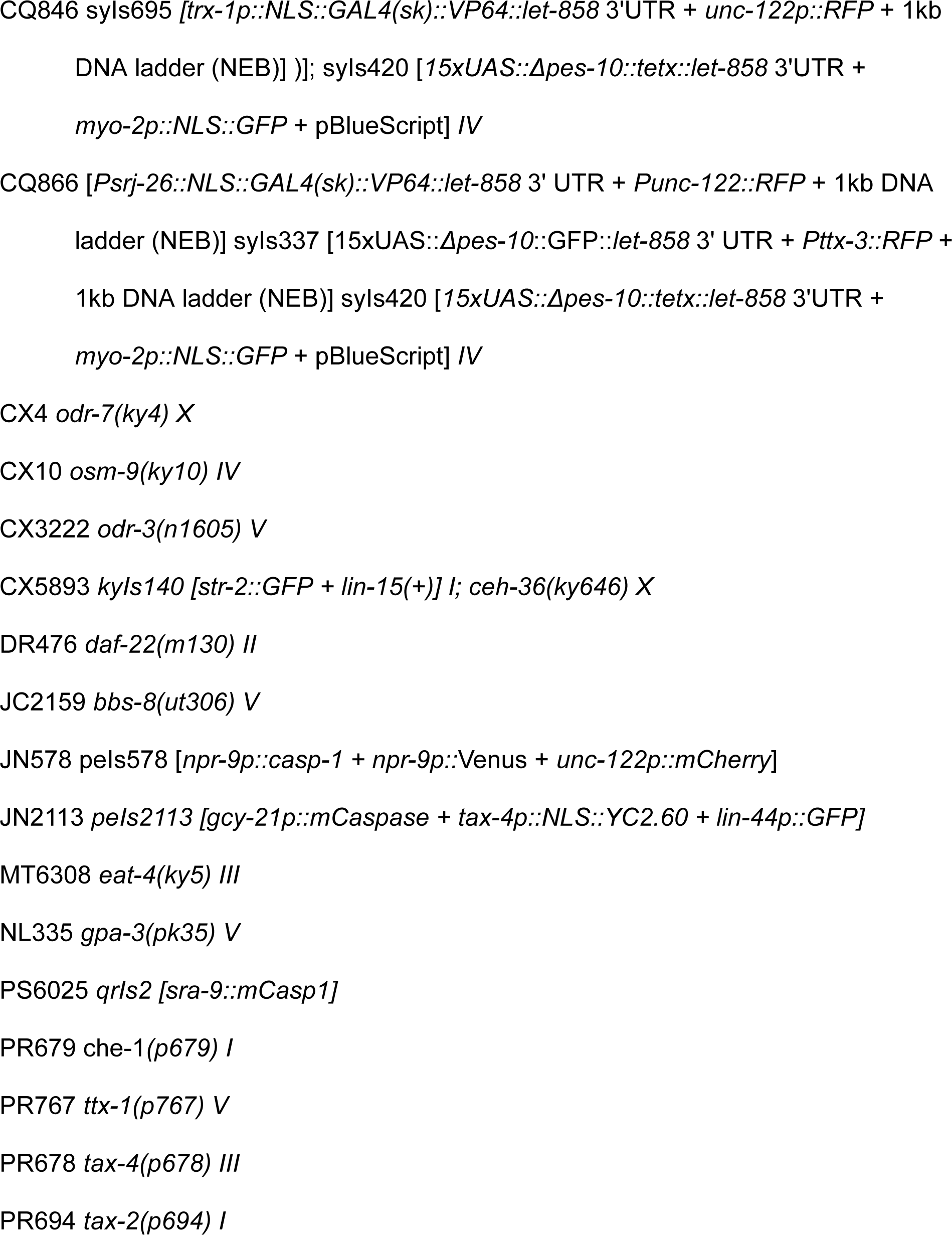
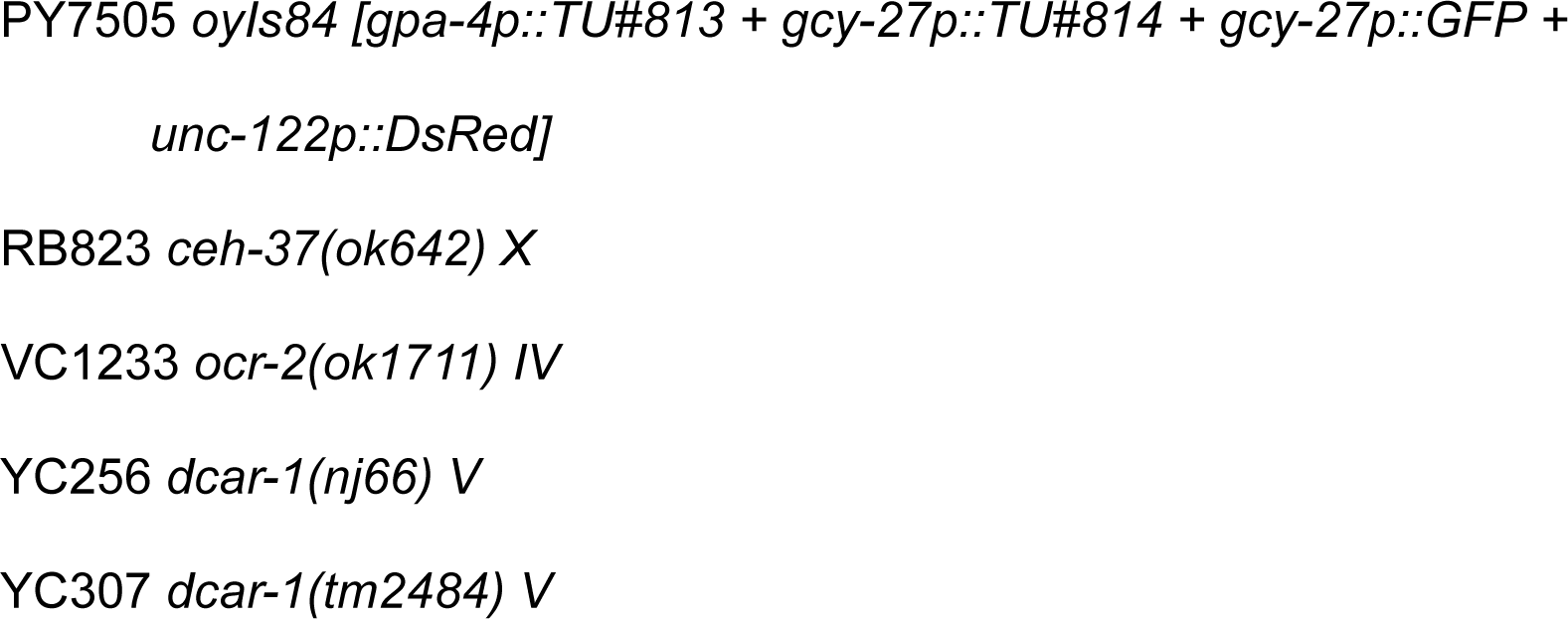

### Preparation of *C. elegans* Lysates

Bleach synchronized worms were raised on HG plates seeded with OP50. Day 1 adults were collected by washing with PBS buffer pH 7.4 [0.137M NaCl, 2.7mM KCl, 8mM Na2HPO4, 2mM KH2PO4] (Invitrogen) and were washed 3x with PBS to remove residual bacteria. Worms were then pelleted by gravity on ice, and excess buffer was removed. Worms were then lysed on ice by Dounce homogenization or gentle sonication using a needle sonicator. The lysate was then centrifuged at 500xg for 10 minutes in a 4°C refrigerated benchtop centrifuge to separate carcasses from lysate, after which the supernatant was collected and filtered with a 0.2μm syringe filter. Protein content of the lysate was used as a proxy for lysate concentration across trials, as determined using a Qubit Protein Assay kit (ThermoFisher). Lysate was either immediately used in experiments or was flash frozen in liquid nitrogen and stored at −80°C for later use.

### Preparation of *E. coli* Lysates

Overnight cultures of OP50 *E. coli* in LB [10 g/L Bacto-tryptone, 5 g/L Bacto-yeast, 10 g/L NaCl] were centrifuged at 2000xg for 5 minutes and were washed 2x with PBS buffer pH 7.4, centrifuging after each wash. Bacterial pellets were resuspended in minimal PBS buffer and were sonicated on ice using a needle sonicator. Unlysed bacteria were pelleted by centrifuging at 2000xg for 5 minutes in a 4°C refrigerated benchtop centrifuge, and the supernatant was filtered through a 0.2μm filter. Protein content of the lysate was used as a proxy for lysate concentration across trials, as determined using a Qubit Protein Assay kit (ThermoFisher). Bacterial lysates were used immediately or were flash frozen in liquid nitrogen and stored at −80°C for future use.

### Preparation of *C. elegans* Conditioned Media

Conditioned media was prepared by washing bleach synchronized day 1 adult DR476 worms from HG plates seeded with OP50 with M9. The worms were washed 1x with M9 and allowed to settle by gravity, and supernatant was removed. 300μL of S Basal [5.85 g/L NaCl, 1 g/L K_2_ HPO_4_, 6 g/L KH_2_PO_4_, 1 ml/L cholesterol (5 mg/ml in ethanol)] was added for every 50μL of worm pellet, and 1.5mL of worm:S Basal solution was transferred into each well of a 6-well plate for 18 hours in a 20°C incubator. Following this incubation, each well was visually confirmed to not contain dead or lysed worms. Worms and the conditioned media were gently transferred to 15mL conical tubes using a wide orifice pipette tip and were allowed to settle by gravity on ice. The supernatant was collected and centrifuged 6500xg for 10 min in a 4°C benchtop centrifuge. The supernatant was collected and passed through a 0.2μm syringe filter. Conditioned media was immediately used for experiments or was flash frozen in liquid nitrogen and stored at −80°C for later use.

### Necrotaxis assay

The day before a necrotaxis assay was to be performed, ≥7 60mm NGM plates were prepared for each condition to be tested. The bottom of each plate was marked with a point directly in the center of the plate and two parallel lines each 1cm from the center point. 20 or 25μL of OP50 culture in LB was deposited on the far side of each line, being careful that the drop did not cross the designated lines. On the day of the assay, 40μL of PBS buffer was applied in a continuous line on each of the negative control plates. For lysate experimental plates, 40μL of PBS buffer was applied along the marked line on one side of the plate, while 40μL of lysate (diluted to 5μg/μL protein concentration with PBS unless stated otherwise) was applied on the line on the opposite side. Each line of PBS or lysate was carefully placed such that there were no breaks in the liquid and damage to the surface of the agar was avoided. 1μL of 6.6% sodium azide in water was then applied to each food patch to paralyze worms and capture their first choice. Bleach synchronized day 1 adult worms were then washed off HG plates seeded with OP50 using M9 buffer and were washed 3x with M9 buffer to remove excess bacteria, settling by gravity between each wash. 4-5μL of worms were applied to the center point of each plate, with an ideal plate having 80-130 worms. The total time between beginning to apply PBS or lysate to plates and placing worms on plates was always less than one hour, and the plates were allowed to completely dry before worms were placed onto them. One hour after worms were placed onto plates, an image of each plate was acquired using a Basler scA1600 17fm camera. The number of worms on the plate, as well as the number of worms that crossed the PBS or lysate lines, was scored using FIJI. The preference index was calculated as: (# worms on lysate side - # worms on buffer control side)/(Total worms on the plate). For negative control PBS vs PBS plates, one side of the plate was arbitrarily determined to be the “lysate” side for this calculation before the experiment was begun. Plates were only used in downstream analysis if ≥30 worms crossed the PBS and/or lysate lines in total, and a technical replicate was only included if at least 4 plates met this criterion. A PBS vs PBS negative control was included in each experiment for each genotype of worm assayed to ensure that if a negative preference index was observed, indicating avoidance, that this effect was not due to incidental environmental factors. In some cases, multiple experiments were performed using the same controls (e.g. PBS vs PBS) as long as both the assay plates were prepared and the worms were all plated at the same time.

Necrotaxis assays were also performed on 35mm plates when screening GPCR mutants (Figure 4A) and HPLC fractions (Figure 7C). In these assays, 8μL of OP50 was used to seed the plates, barriers of 10μL of buffer/lysate/HPLC fractions were used, and no sodium azide was applied to the plate. Plates were imaged 30 minutes after worms were placed onto them.

### Lysate Dialysis

*C. elegans* lysates were dialyzed in 2kDa or 20kDa MWCO Slidealyzer™ cassettes, or in 50kDa dialysis tubing (Spectrum Labs). Samples were placed in >1000x volume of PBS pH7.4 overnight at 4°C with constant stirring and were used immediately in necrotaxis assays after retrieval.

### Lysate lipid depletion

Lipid depletion was performed using Cleanascite™ lipid depletion reagent (Biotech Support Group), which depletes lipids from samples to a similar extent as chloroform extraction (Castro et al., 2000). The Cleanascite reagent was centrifuged at 3000xg for 15 minutes at 4°C in a benchtop refrigerated centrifuge. The supernatant was discarded and the reagent was then washed 1x in PBS followed by centrifugation. The washed Cleanascite reagent was then added in a 1:1 ratio to PBS or *C. elegans* lysate (standardized to 5μg/μL protein concentration in PBS). The PBS:Cleanascite, Lysate:Cleanascite, or control lysate samples were incubated at 4°C on a rotator to ensure constant gentle mixing. 18-20 hours later, the samples were centrifuged 3000xg for 15 minutes at 4°C, and the supernatant of each was used in necrotaxis assays. In necrotaxis assays, PBS:Cleanascite supernatant was used as the control on Lysate:Cleanascite plates to control for any residual components of the Cleanascite buffer or beads that might affect the worms’ behavior.

### Lysate glycan depletion

Glycan depletion was performed using wheat germ agglutinin (WGA) coated beads (Vector Laboratories). WGA beads were washed 4x with PBS, centrifuging at 2000xg for 1 minute at 4°C between each wash. WGA beads were added to Lysate at a 1:2 ratio to PBS or *C. elegans* lysate (standardized to 5μg/μL protein concentration in PBS). The PBS:WGA, Lysate:WGA, or control lysate samples were incubated at 4°C on a rotator to ensure constant gentle mixing. 18-20 hours later, the samples were centrifuged at 2000xg for 1 minute at 4°C and the supernatant was used in necrotaxis assays. In necrotaxis assays, PBS:WGA supernatant was used as the control on Lysate:WGA plates instead of PBS alone to control for any residual components of the WGA beads or buffer that might affect the worms’ behavior.

### *C. elegans* RNA extraction

Bleach synchronized day 1 adult N2 *C. elegans* were washed off of HG plates seeded with OP50 using M9 buffer and were allowed to pellet by gravity. TRIzol™ (Invitrogen) was added to the worms at a 9:1 ratio, and the suspension was frozen at −80°C. RNA was extracted from the worms using a *mir*Vana™ miRNA Isolation kit (Invitrogen) per the kit’s instructions. sRNA and large RNA populations were eluted with nuclease free water, and the nucleic acid concentration was determined by nanodrop. RNAs were stored at −20°C until they were used in necrotaxis assays.

### *C. elegans* DNA extraction

Bleach synchronized day 1 adult N2 *C. elegans* were washed off of HG plates seeded with OP50 using M9 buffer and were allowed to pellet by gravity. The supernatant was removed, and the worm pellet was aliquoted into 250μL volumes and frozen at −20°C. The samples were then thawed and 750μL of lysis buffer [200mM NaCl, 100mM Tris-HCl pH8.5, 50mM EDTA pH8.0, 0.5% SDS, 0.1mg/mL proteinase K] was added to each aliquot. The worms were incubated at 65°C in a Thermomixer for 1 hour with vortexing every 15 minutes, and then the proteinase K was heat-inactivated at 95°C for 20 minutes in a Thermomixer. RNAseA (Omega Biotek) was then added to each sample to 0.1mg/mL and samples were incubated at 37°C for 1 hour in a Thermomixer. 250μL of phenol:chloroform:isoamylic acid was added to each sample, and each sample was shaken vigorously and centrifuged at 13000rpm at room temperature for 15 minutes. 400μL of the top (aqueous) phase was transferred to a new tube and 40μL of 3M sodium acetate and 1200μL of 100% ethanol were added to each tube. The samples were left at −80°C overnight to precipitate DNA. The samples were then centrifuged at 13000rpm for 15 minutes at 4°C, and the supernatant was discarded. The pellet was washed once with ice cold 75% ethanol, and then was centrifuged again 13000rpm for 15 minutes at 4°C. All supernatant was discarded and the pellets were air dried. Each DNA pellet was resuspended in 50μL of nuclease free water and the DNA concentration was determined using a Nanodrop. DNA was stored at −20°C until used in a necrotaxis assay.

### Trichloroacetic acid protein precipitation

100% trichloroacetic acid was added to *C. elegans* or OP50 lysate (protein concentration 10μg/μL) to a final concentration of 20% and the sample was incubated on ice for 30 minutes. The lysate was then centrifuged at 5000xg at 4°C in a refrigerated benchtop centrifuge for 1 minute. The supernatant was removed, and the pellet was washed twice with 90% acetone 10mM HCl. The supernatant was removed, and the pellet was dried to remove residual acetone. The pellet was resuspended in 1.5x of its original volume in PBS pH7.4. The solution was then sonicated using a needle sonicator to disrupt the pellet. Insoluble aggregate was then pelleted by centrifuging at 21000xg for 1 minute, and the supernatant was transferred to a new tube and used immediately in experiments.

### Acetone Protein Precipitation

100% acetone (prechilled to −20°C) was added to *C. elegans* lysate to a final concentration of 80% with gentle mixing and samples were incubated at −20°C for 60 minutes. Samples were then centrifuged at 6000xg 4°C and the supernatant was discarded. Protein pellets were air dried to remove residual acetone and then were resuspended in PBS pH7.4. The samples were rehydrated by incubating them on ice for one hour followed by gentle disruption with a needle sonicator. The protein was then centrifuged at 21000xg 4°C for 10 minutes to pellet residual insoluble protein. The supernatant was transferred to a new tube and the protein concentration of the resolubilized protein was determined using a Qubit Protein Quantification kit. All samples were immediately used in downstream experiments.

### FPLC Size Fractionation

*C. elegans* protein precipitated using acetone (see above) and resuspended in PBS or ddH_2_O was centrifuged for 5 minutes at 4°C 12500xg in 0.22μm spin columns (Costar). Size exclusion chromatography was performed at 4°C at a flowrate of 0.4 mL/min on an ÄKTA Fast Performance Liquid Chromatography (GE Healthcare) system using a BIO-RAD ENrich SEC 70 10 x 300 column (#7801070) with either Phosphate-Buffered Saline (PBS) or water as the eluent. Pure fractions were used for downstream analysis and functional assays.

### HPLC Reverse Phase Fractionation

Without concentrating, samples derived from FPLC fractionation were further purified using semi-prep HPLC on a Synergi 4μm Hydro-RP 80Å LC 250x4.6mm column. Solvent A was 0.2% formic acid in MeCN and Solvent B was 0.1% formic acid in H_2_O. The following gradient was applied during elution: 100% B to 5% A in B (0-10 min), 5% A in B to 10% A in B (10-12 min), 10%A in B to 30% A in B (12-16 min), 30% A in B to 100% A (16-20min) at a flow rate of 1mL/minute. 100μL of sample was injected for each HPLC elution. Samples of the same fractions from different injections were combined and concentrated using a speedvac.

### Proteinase K Digestion of Precipitated and Resolubilized Protein

*C. elegans* acetone precipitated protein (see above; 1mg/mL protein concentration) was incubated at 56°C with 0.2mg/mL Proteinase K (Invitrogen) for 1 hour. Mock controls were performed by incubating lysate at 56°C without Proteinase K, and negative control lysate was incubated on ice without Proteinase K. All samples were immediately used in subsequent assays.

### NMR Sample Preparation

The combined Fraction #3 from HPLC of UV Peak #2 of ∼60 HPLC runs was dried using speedvac. The sample was dissolved in 200μL D_2_O for NMR analysis.

### Proteinase K Digestion of Lysate Preceding Protein Precipitation

*C. elegans* lysate (5mg/mL protein concentration) was incubated at 56°C with 0.2mg/mL Proteinase K (Invitrogen) for 1 hour. Mock controls were performed by incubating lysate at 56°C without Proteinase K, and negative control lysate was incubated on ice without Proteinase K. Protein in each condition was then precipitated using acetone (see above) and pellets were resolubilized in a volume of PBS equal to the starting amount of lysate. All samples were immediately used in subsequent assays.

### Statistical analysis

All statistics were performed using R (v4.4.2). Data wrangling was performed using the reshape2 (v1.4.4), DescTools (v0.99.53), and tidyverse (v2.0.0) packages. Specific statistical tests used are denoted in figure legends.

## Acknowledgements

We thank the members of the Murphy lab for feedback on this manuscript. Funding for this work was provided by the Simons Foundation Collaboration on Plasticity in the Aging Brain (SCPAB) to CTM, an NIH Director’s Pioneer Award to CTM (DP1AG077430), and the Glenn Foundation for Medical Research. ET is a recipient of the Life Sciences Research Foundation (LSRF) fellowship supported by the Simons Foundation Autism Research Initiative (SFARI). EL is a recipient of an EMBO Postdoctoral Fellowship. JZH is a recipient of the Life Sciences Research Foundation (LSRF) fellowship supported by the Open Philanthropy Project. Some strains were provided by the *Caenorhabditis* Genetics Center (CGC), which is funded by NIH Office of Research Infrastructure Programs (P40 OD010440). We also thank the Princeton Mass Spectrometry and NMR cores for their assistance in this work.

## Literature Cited

(1) Chakraborty, T. S.; Gendron, C. M.; Lyu, Y.; Munneke, A. S.; DeMarco, M. N.; Hoisington, Z. W.; Pletcher, S. D. Sensory Perception of Dead Conspecifics Induces Aversive Cues and Modulates Lifespan through Serotonin in Drosophila. Nat Commun 2019, 10, 2365. 10.1038/s41467-019-10285-y.

(2) Choe, D.-H.; Millar, J. G.; Rust, M. K. Chemical Signals Associated with Life Inhibit Necrophoresis in Argentine Ants. Proceedings of the National Academy of Sciences 2009, 106 (20), 8251–8255. 10.1073/pnas.0901270106.

(3) Frisch, K. V. Zur Psychologie Des Fisch-Schwarmes. [Psychology of Preference in Fish.]. Naturwissenschaften 1938, 26, 601–606. 10.1007/BF01590598.

(4) Masuda, M.; Ihara, S.; Mori, N.; Koide, T.; Miyasaka, N.; Wakisaka, N.; Yoshikawa, K.; Watanabe, H.; Touhara, K.; Yoshihara, Y. Identification of Olfactory Alarm Substances in Zebrafish. Current Biology 2024, 34 (7), 1377–1389.e7. 10.1016/j.cub.2024.02.003.

(5) Mathuru, A. S.; Kibat, C.; Cheong, W. F.; Shui, G.; Wenk, M. R.; Friedrich, R. W.; Jesuthasan, S. Chondroitin Fragments Are Odorants That Trigger Fear Behavior in Fish. Current Biology 2012, 22 (6), 538–544. 10.1016/j.cub.2012.01.061.

(6) Wisenden, B. D.; Millard, M. C. Aquatic Flatworms Use Chemical Cues from Injured Conspecifics to Assess Predation Risk and to Associate Risk with Novel Cues. Animal Behaviour 2001, 62 (4), 761–766. 10.1006/anbe.2001.1797.

(7) Zhou, Y.; Loeza-Cabrera, M.; Liu, Z.; Aleman-Meza, B.; Nguyen, J. K.; Jung, S.-K.; Choi, Y.; Shou, Q.; Butcher, R. A.; Zhong, W. Potential Nematode Alarm Pheromone Induces Acute Avoidance in Caenorhabditis Elegans. Genetics 2017, 206 (3), 1469–1478. 10.1534/genetics.116.197293.

(8) Hernandez-Lima, M. A.; Seo, B.; Urban, N. D.; Truttmann, M. C. Modulation of *C. Elegans* Behavior, Fitness, and Lifespan by AWB/ASH-Dependent Death Perception. Current Biology 2025. 10.1016/j.cub.2025.03.071.

(9) Debru, C. A Particular Form of Chemotaxis: Necrotaxis. An Historical View. Blood Cells 1993, 19 (1), 5–19; discussion 20-23.

(10) McDonald, B.; Pittman, K.; Menezes, G. B.; Hirota, S. A.; Slaba, I.; Waterhouse, C. C. M.; Beck, P. L.; Muruve, D. A.; Kubes, P. Intravascular Danger Signals Guide Neutrophils to Sites of Sterile Inflammation. Science 2010, 330 (6002), 362–366. 10.1126/science.1195491.

(11) Bessis, M.; Burté, B. Positive and Negative Chemotaxis as Observed after the Destruction of a Cell by U. V. or Laser Microbeams. Tex Rep Biol Med 1965, 23, Suppl 1:204–12.

(12) Ragot, R. Negative Necrotaxis. Blood Cells 1993, 19 (1), 81–88; discussion 88-90.

(13) Bargmann, C. I. Chemosensation in C. Elegans. In WormBook: The Online Review of C. elegans Biology *[Internet]*; WormBook, 2006.

(14) Hernandez-Lima, M. A.; Seo, B.; Urban, N. D.; Truttmann, M. C. C. Elegans Behavior, Fitness, and Lifespan, Are Modulated by AWB/ASH-Dependent Death Perception. bioRxiv October 9, 2024, p 2024.10.07.617097. 10.1101/2024.10.07.617097.

(15) Macosko, E. Z.; Pokala, N.; Feinberg, E. H.; Chalasani, S. H.; Butcher, R. A.; Clardy, J.; Bargmann, C. I. A Hub-and-Spoke Circuit Drives Pheromone Attraction and Social Behaviour in C. Elegans. Nature 2009, 458 (7242), 1171–1175. 10.1038/nature07886.

(16) Lanjuin, A.; VanHoven, M. K.; Bargmann, C. I.; Thompson, J. K.; Sengupta, P. Otx/Otd Homeobox Genes Specify Distinct Sensory Neuron Identities in C. Elegans. Developmental Cell 2003, 5 (4), 621–633. 10.1016/S1534-5807(03)00293-4.

(17) Blacque, O. E.; Reardon, M. J.; Li, C.; McCarthy, J.; Mahjoub, M. R.; Ansley, S. J.; Badano, J. L.; Mah, A. K.; Beales, P. L.; Davidson, W. S.; Johnsen, R. C.; Audeh, M.; Plasterk, R. H. A.; Baillie, D. L.; Katsanis, N.; Quarmby, L. M.; Wicks, S. R.; Leroux, M. R. Loss of C. Elegans BBS-7 and BBS-8 Protein Function Results in Cilia Defects and Compromised Intraflagellar Transport. Genes Dev 2004, 18 (13), 1630–1642. 10.1101/gad.1194004.

(18) Fujiwara, M.; Ishihara, T.; Katsura, I. A Novel WD40 Protein, CHE-2, Acts Cell-Autonomously in the Formation of C. Elegans Sensory Cilia. Development 1999, 126 (21), 4839–4848. 10.1242/dev.126.21.4839.

(19) Altun, Z. F.; Herndon, L. A.; Wolkow, C. A.; Crocker, C.; Lints, R.; Hall, D. H. WormAtlas, 2002. http://www.wormatlas.org.

(20) Chang, S.; Johnston, R. J.; Hobert, O. A Transcriptional Regulatory Cascade That Controls Left/Right Asymmetry in Chemosensory Neurons of C. Elegans. Genes Dev. 2003, 17 (17), 2123–2137. 10.1101/gad.1117903.

(21) Koga, M.; Ohshima, Y. The *C. Elegans Ceh*-*36* Gene Encodes a Putative Homemodomain Transcription Factor Involved in Chemosensory Functions of ASE and AWC Neurons. Journal of Molecular Biology 2004, 336 (3), 579–587. 10.1016/j.jmb.2003.12.037.

(22) Satterlee, J. S.; Sasakura, H.; Kuhara, A.; Berkeley, M.; Mori, I.; Sengupta, P. Specification of Thermosensory Neuron Fate in C. Elegans Requires Ttx-1, a Homolog of Otd/Otx. Neuron 2001, 31 (6), 943–956. 10.1016/S0896-6273(01)00431-7.

(23) Sengupta, P.; Colbert, H. A.; Bargmann, C. I. The C. Elegans Gene Odr-7 Encodes an Olfactory-Specific Member of the Nuclear Receptor Superfamily. Cell 1994, 79 (6), 971–980. 10.1016/0092-8674(94)90028-0.

(24) Uchida, O.; Nakano, H.; Koga, M.; Ohshima, Y. The C. Elegans Che-1 Gene Encodes a Zinc Finger Transcription Factor Required for Specification of the ASE Chemosensory Neurons. Development 2003, 130 (7), 1215–1224. 10.1242/dev.00341.

(25) Wang, H.; Liu, J.; Gharib, S.; Chai, C. M.; Schwarz, E. M.; Pokala, N.; Sternberg, P. W. cGAL, a Temperature-Robust GAL4–UAS System for Caenorhabditis Elegans. Nat Methods 2017, 14 (2), 145–148. 10.1038/nmeth.4109.

(26) Philbrook, A.; O’Donnell, M. P.; Grunenkovaite, L.; Sengupta, P. Cilia Structure and Intraflagellar Transport Differentially Regulate Sensory Response Dynamics within and between C. Elegans Chemosensory Neurons. PLOS Biology 2024, 22 (11), e3002892. 10.1371/journal.pbio.3002892.

(27) Zhang, X.; Liu, J.; Pan, T.; Ward, A.; Liu, J.; Xu, X. Z. S. A Cilia-Independent Function of BBSome Mediated by DLK MAPK Signaling in C. Elegans Photosensation. Dev Cell 2022, 57 (12), 1545–1557.e4. 10.1016/j.devcel.2022.05.005.

(28) Krzyzanowski, M. C.; Woldemariam, S.; Wood, J. F.; Chaubey, A. H.; Brueggemann, C.; Bowitch, A.; Bethke, M.; L’Etoile, N. D.; Ferkey, D. M. Aversive Behavior in the Nematode C. Elegans Is Modulated by cGMP and a Neuronal Gap Junction Network. PLOS Genetics 2016, 12 (7), e1006153. 10.1371/journal.pgen.1006153.

(29) Coburn, C. M.; Bargmann, C. I. A Putative Cyclic Nucleotide–Gated Channel Is Required for Sensory Development and Function in C. Elegans. Neuron 1996, 17 (4), 695–706. 10.1016/S0896-6273(00)80201-9.

(30) Komatsu, H.; Mori, I.; Rhee, J. S.; Akaike, N.; Ohshima, Y. Mutations in a Cyclic Nucleotide-Gated Channel Lead to Abnormal Thermosensation and Chemosensation in C. Elegans. Neuron 1996, 17 (4), 707–718. 10.1016/s0896-6273(00)80202-0.

(31) Colbert, H. A.; Smith, T. L.; Bargmann, C. I. OSM-9, A Novel Protein with Structural Similarity to Channels, Is Required for Olfaction, Mechanosensation, and Olfactory Adaptation inCaenorhabditis Elegans. J. Neurosci. 1997, 17 (21), 8259–8269. 10.1523/JNEUROSCI.17-21-08259.1997.

(32) St Ange, J.; Weng, Y.; Kaletsky, R.; Stevenson, M. E.; Moore, R. S.; Zhou, S.; Murphy, C. T. Adult Single-Nucleus Neuronal Transcriptomes of Insulin Signaling Mutants Reveal Regulators of Behavior and Learning. Cell Genom 2024, 4 (12), 100720. 10.1016/j.xgen.2024.100720.

(33) Pu, L.; Wang, J.; Lu, Q.; Nilsson, L.; Philbrook, A.; Pandey, A.; Zhao, L.; Schendel, R. van; Koh, A.; Peres, T. V.; Hashi, W. H.; Myint, S. L.; Williams, C.; Gilthorpe, J. D.; Wai, S. N.; Brown, A.; Tijsterman, M.; Sengupta, P.; Henriksson, J.; Chen, C. Dissecting the Genetic Landscape of GPCR Signaling through Phenotypic Profiling in C. Elegans. Nat Commun 2023, 14 (1), 8410. 10.1038/s41467-023-44177-z.

(34) Taylor, S. R.; Santpere, G.; Weinreb, A.; Barrett, A.; Reilly, M. B.; Xu, C.; Varol, E.; Oikonomou, P.; Glenwinkel, L.; McWhirter, R.; Poff, A.; Basavaraju, M.; Rafi, I.; Yemini, E.; Cook, S. J.; Abrams, A.; Vidal, B.; Cros, C.; Tavazoie, S.; Sestan, N.; Hammarlund, M.; Hobert, O.; Miller, D. M. Molecular Topography of an Entire Nervous System. Cell 2021, 184 (16), 4329–4347.e23. 10.1016/j.cell.2021.06.023.

(35) Bono, M. de; Maricq, A. V. NEURONAL SUBSTRATES OF COMPLEX BEHAVIORS IN C. ELEGANS. Annual Review of Neuroscience 2005, 28 (Volume 28, 2005), 451–501. 10.1146/annurev.neuro.27.070203.144259.

(36) Hilliard, M. A.; Bargmann, C. I.; Bazzicalupo, P. *C. Elegans* Responds to Chemical Repellents by Integrating Sensory Inputs from the Head and the Tail. Current Biology 2002, 12 (9), 730–734. 10.1016/S0960-9822(02)00813-8.

(37) Sambongi, Y.; Nagae, T.; Liu, Y.; Yoshimizu, T.; Takeda, K.; Wada, Y.; Futai, M. Sensing of Cadmium and Copper Ions by Externally Exposed ADL, ASE, and ASH Neurons Elicits Avoidance Response in Caenorhabditis Elegans. NeuroReport 1999, 10 (4), 753.

(38) Ferkey, D. M.; Sengupta, P.; L’Etoile, N. D. Chemosensory Signal Transduction in Caenorhabditis Elegans. Genetics 2021, 217 (3), iyab004. 10.1093/genetics/iyab004.

(39) Hart, A. C.; Sims, S.; Kaplan, J. M. Synaptic Code for Sensory Modalities Revealed by C. Elegans GLR-1 Glutamate Receptor. Nature 1995, 378 (6552), 82–85. 10.1038/378082a0.

(40) Lee, R. Y.; Sawin, E. R.; Chalfie, M.; Horvitz, H. R.; Avery, L. EAT-4, a Homolog of a Mammalian Sodium-Dependent Inorganic Phosphate Cotransporter, Is Necessary for Glutamatergic Neurotransmission in Caenorhabditis Elegans. J Neurosci 1999, 19 (1), 159–167. 10.1523/JNEUROSCI.19-01-00159.1999.

(41) Sweeney, S. T.; Broadie, K.; Keane, J.; Niemann, H.; O’Kane, C. J. Targeted Expression of Tetanus Toxin Light Chain in Drosophila Specifically Eliminates Synaptic Transmission and Causes Behavioral Defects. Neuron 1995, 14 (2), 341–351. 10.1016/0896-6273(95)90290-2.

(42) Piggott, B. J.; Liu, J.; Feng, Z.; Wescott, S. A.; Xu, X. Z. S. The Neural Circuits and Synaptic Mechanisms Underlying Motor Initiation in C. Elegans. Cell 2011, 147 (4), 922–933. 10.1016/j.cell.2011.08.053.

(43) Kahn-Kirby, A. H.; Dantzker, J. L. M.; Apicella, A. J.; Schafer, W. R.; Browse, J.; Bargmann, C. I.; Watts, J. L. Specific Polyunsaturated Fatty Acids Drive TRPV-Dependent Sensory Signaling In Vivo. Cell 2004, 119 (6), 889–900. 10.1016/j.cell.2004.11.005.

(44) Zugasti, O.; Bose, N.; Squiban, B.; Belougne, J.; Kurz, C. L.; Schroeder, F. C.; Pujol, N.; Ewbank, J. J. Activation of a G Protein–Coupled Receptor by Its Endogenous Ligand Triggers the Innate Immune Response of Caenorhabditis Elegans. Nat Immunol 2014, 15 (9), 833–838. 10.1038/ni.2957.

(45) Aoki, R.; Yagami, T.; Sasakura, H.; Ogura, K.; Kajihara, Y.; Ibi, M.; Miyamae, T.; Nakamura, F.; Asakura, T.; Kanai, Y.; Misu, Y.; Iino, Y.; Ezcurra, M.; Schafer, W. R.; Mori, I.; Goshima, Y. A Seven-Transmembrane Receptor That Mediates Avoidance Response to Dihydrocaffeic Acid, a Water-Soluble Repellent in Caenorhabditis Elegans. J. Neurosci. 2011, 31 (46), 16603–16610. 10.1523/JNEUROSCI.4018-11.2011.

(46) Koontz, L. Chapter One - TCA Precipitation. In Methods in Enzymology; Lorsch, J., Ed.; Laboratory Methods in Enzymology: Protein Part C; Academic Press, 2014; Vol. 541, pp 3–10. 10.1016/B978-0-12-420119-4.00001-X.

(47) Baghalabadi, V.; Doucette, A. A. Mass Spectrometry Profiling of Low Molecular Weight Proteins and Peptides Isolated by Acetone Precipitation. Analytica Chimica Acta 2020, 1138, 38–48. 10.1016/j.aca.2020.08.057.

(48) He, M.; Zhou, X.; Wang, X. Glycosylation: Mechanisms, Biological Functions and Clinical Implications. Sig Transduct Target Ther 2024, 9 (1), 194. 10.1038/s41392-024-01886-1.

(49) Butcher, R. A.; Ragains, J. R.; Li, W.; Ruvkun, G.; Clardy, J.; Mak, H. Y. Biosynthesis of the Caenorhabditis Elegans Dauer Pheromone. Proceedings of the National Academy of Sciences 2009, 106 (6), 1875–1879. 10.1073/pnas.0810338106.

(50) Le, H. H.; Wrobel, C. J.; Cohen, S. M.; Yu, J.; Park, H.; Helf, M. J.; Curtis, B. J.; Kruempel, J. C.; Rodrigues, P. R.; Hu, P. J.; Sternberg, P. W.; Schroeder, F. C. Modular Metabolite Assembly in Caenorhabditis Elegans Depends on Carboxylesterases and Formation of Lysosome-Related Organelles. eLife 2020, 9, e61886. 10.7554/eLife.61886.

(51) Kaletsky, R.; Yao, V.; Williams, A.; Runnels, A. M.; Tadych, A.; Zhou, S.; Troyanskaya, O. G.; Murphy, C. T. Transcriptome Analysis of Adult Caenorhabditis Elegans Cells Reveals Tissue-Specific Gene and Isoform Expression. PLoS Genet 2018, 14 (8), e1007559. 10.1371/journal.pgen.1007559.

(52) Quach, K. T.; Chalasani, S. H. Flexible Reprogramming of *Pristionchus Pacificus* Motivation for Attacking *Caenorhabditis Elegans* in Predator-Prey Competition. Current Biology 2022, 32 (8), 1675–1688.e7. 10.1016/j.cub.2022.02.033.

